# Untargeted Mass Spectrometry-Based Metabolomics Tracks Molecular Changes in Raw and Processed Foods and Beverages

**DOI:** 10.1101/347716

**Authors:** Julia M. Gauglitz, Christine M. Aceves, Alexander A. Aksenov, Gajender Aleti, Jehad Almaliti, Amina Bouslimani, Elizabeth A. Brown, Anaamika Campeau, Andrés Mauricio Caraballo-Rodríguez, Rama Chaar, Ricardo R. da Silva, Alyssa M. Demko, Francesca Di Ottavio, Emmanuel Elijah, Madeleine Ernst, L. Paige Ferguson, Xavier Holmes, Justin J.J. van der Hooft, Alan K. Jarmusch, Lingjing Jiang, Kyo Bin Kang, Irina Koester, Brian Kwan, Bohan Ni, Jie Li, Yueying Li, Alexey V. Melnik, Carlos Molina-Santiago, Aaron L. Oom, Morgan W. Panitchpakdi, Daniel Petras, Robert Quinn, Nicole Sikora, Katharina Spengler, Bahar Teke, Anupriya Tripathi, Sabah Ul-Hasan, Fernando Vargas, Alison Vrbanac, Anthony Q. Vu, Steven C Wang, Kelly Weldon, Kayla Wilson, Jacob M. Wozniak, Michael Yoon, Nuno Bandeira, Pieter C. Dorrestein

**Affiliations:** Collaborative Mass Spectrometry Innovation Center, Skaggs School of Pharmacy and Pharmaceutical Sciences, University of California, San Diego.; Skaggs School of Pharmacy and Pharmaceutical Sciences, University of California, San Diego; Center for Microbiome Innovation, University of California, San Diego.; Mammalian Genomics, J. Craig Venter Institute, San Diego.; Scripps Institution of Oceanography, University of California, San Diego; Department of Pharmaceutical Sciences, Faculty of Pharmacy, The University of Jordan, Amman, 11942, Jordan; Department of Pharmacology, University of California, San Diego.; University of Teramo, Faculty of Bioscience, Te, Italy; Bioinformatics Group, Wageningen University, Wageningen, The Netherlands; Department of Family Medicine and Public Health, University of California, San Diego.; Departamento de Microbiologfa, Instituto de Hortofruticultura Subtropical y Mediterranea “La Mayora”, Universidad de Málaga, Bulevar Louis Pasteur 31 (Campus Universitario de Teatinos), 29071 Málaga, Spain.; Department of Medicine, University of California, San Diego; School of Natural Sciences, University of California Merced, Merced, CA, 95343, USA; Division of Biological Sciences, University of California at San Diego, La Jolla, CA; Biomedical Sciences Graduate Program, University of California San Diego, La Jolla, CA, USA; Department of Computer Science and Engineering, University of California, San Diego.; Departments of Pharmacology and Pediatrics, University of California, San Diego.

**Author notes:** **Author Contributions:** PCD and NB conceptualized the idea. CMA, EAB, AAA, AMD, AT, AMCR, AM, CMS, FDO, KBK, KS, IK, MWP, BK, AC, DP, SCW, JMG, KAW, JMW, BN, BT, JA, RA, AV, YL collected the samples and metadata JMG, FDO, MWP, AKJ collected the MS data JMG processed the data JMG, AMCR, KBK, ALO, DP, AC performed molecular networking. ME created heatmaps. RRdS, ME, AKJ, AT, JMG performed preprocessing of data and statistical analysis. JMG, AKJ, JVDH, ME, RRdS, AAA, AM, AT, DP, NL, AC, JMW, PCD, JA, IK, AV, FV, RAQ, AB, MWP, FDO, GA, ALO analyzed the data. JMG, AKJ, JVDH, SCW, AMCR, AAA, AM, AMD, KBK, IK, ME, ALO, BK, DP, FV, AC, AB, MWP, FDO, NB, JMW PCD, authored the manuscript.

**Keywords:** Untargeted mass spectrometry, metabolomics, Food, Tea, Coffee, Tomato, Milk, Yogurt, LC-MS/MS

## Abstract

A major aspect of our daily lives is the need to acquire, store and prepare our food. Storage and preparation can have drastic effects on the compositional chemistry of our foods, but we have a limited understanding of the temporal nature of processes such as storage, spoilage, fermentation and brewing on the chemistry of the foods we eat. Here, we performed a temporal analysis of the chemical changes in foods during common household preparations using untargeted mass spectrometry and novel data analysis approaches. Common treatments of foods such as home fermentation of yogurt, brewing of tea, spoilage of meats and ripening of tomatoes altered the chemical makeup through time, through both chemical and biological processes. For example, brewing tea altered its composition by increasing the diversity of molecules, but this change was halted after 4 min of brewing. The results indicate that this is largely due to differential extraction of the material from the tea and not modification of the molecules during the brewing process. This is in contrast to the preparation of yogurt from milk, spoilage of meat and the ripening of tomatoes where biological transformations directly altered the foods molecular composition. Comprehensive assessment of chemical changes using multivariate statistics showed the varied impacts of the different food treatments, while analysis of individual chemical changes show specific alterations of chemical families in the different food types. The methods developed here represent novel approaches to studying the changes in food chemistry that can reveal global alterations in chemical profiles and specific transformations at the chemical level.

Highlights
1. We created a reference data set for tomato, milk to yogurt, tea, coffee, turkey and beef.
2. We show that normal preparation and handling affects the molecular make-up.
3. Tea preparation is largely driven by differential extraction.
4. Formation of yogurt involves chemical transformations.
5. The majority of meat molecules are not altered in 5 days at room temperature.

## Introduction

We consume a variety of foods and beverages during any given day, such as fruits, vegetables, dairy products and meats. Food is stored and processed in many different ways before consumption, yet we know very little about the molecular impacts of such “normal” food treatments before we consume them. There is a significant interest and awareness in the population about the molecular contents of food. Consistent with this interest, there are >37,000 articles in Pubmed using the terms “food” and “mass spectrometry” but only ~250 when using the search terms “food”, “untargeted”, and “mass spectrometry” or “metabolomics”. Although as many as 25,000 food molecules are known, the majority of food mass spectrometry studies focus on the detection of insecticides, pesticides and toxins or particular compound classes such as polyphenols to which healthy properties are attributed (Casida & Durkin, 2017, Giorio et al., 2017, Scalbert et al., 2014) and are used to compare different food supplements such as the coffee leaves (Souard, et al., 2018). As a consequence, much of the work is done by targeted methods and/or GC-MS for untargeted methods. Nevertheless, the importance of mass spectrometry as the most sensitive and selective tool currently available to decipher our food is is only expected to grow (Yoshimura, Goto-Inoue, Moriyama, & Zaima, 2016) in areas such as food monitoring during processing (Marshall et al., 2017), especially as the cost per data volume of mass spectrometry has decreased by two orders of magnitude in the past 15 years and is expected to continue to go down (Aksenov et al., 2017). An untargeted approach using LC-MS/MS has not been as widely used to analyze food types and effects of storage and processing and never in conjunction with emerging untargeted mass spectrometry analysis approaches such as mass spectral molecular networking to assess changes based on processing.

Mass spectral molecular networking enables a broad overview of the molecular information, that can be inferred from the MS/MS data (Watrous et al., 2012). For example, molecular networking has been used in food analysis to study Siberian ginseng (Ge, Zhu, Yoshimatsu, & Komatsu 2017) and enabled the characterization of large number of triterpene saponins (Ge, Zhu, Yoshimatsu, & Komatsu 2017). In molecular networking, all identical MS/MS spectra are merged giving a list of unique MS/MS spectra (Watrous et al., 2012). These are then subjected to spectral alignment allowing for spectral matching with offsets based on the parent mass differences. In addition, the MS/MS spectra are putatively annotated against reference spectra within the Global Natural Product Social Molecular Networking (GNPS) platform (Yang et al., 2013, Wang et al., 2016). Matches against the reference libraries constitute level 2 or 3 annotations according to the 2007 metabolomics standards initiative (Sumner et al., 2007). The reference libraries that can be searched against, as their spectra are publicly available or available for purchase (and can be used as private spectral libraries in GNPS), include NIST17, Massbank Europe and North America, ReSpect, CASMI, EMBL metabolomics library, HMDB, and from GNPS contributed MS/MS spectra (Wang M et al., 2016, Aksenov, da Silva, Knight, Lopes, & Dorrestein, 2017, Blaženović, Kind, Ji, & Fiehn, 2018). The resulting molecular networks visualize chemical relationships of detected compounds and provide a powerful tool for in-depth interpretation of chemical transformations.

With these and other widely used mass spectrometry approaches, such as multivariate statistics, we set out to investigate how the molecular make-up of foods is impacted by normal handling before consumption, during processing and preparation. *We hypothesize that many of our methods of sourcing, handling and/or processing of food impact the molecular make-up of the foods and that untargeted mass spectrometry combined with advanced analysis tools can give us insight into these impacts on a molecular level*. Building upon this hypothesis we aim to address some of the following specific questions: 1) How does the molecular composition of a food change based on how it was ripened off the vine or its particular sourcing (tomato), 2) how does improper storage affect the molecular make-up (meat), 3) what are the molecular impacts of starter culture for the preparation of yogurts, 4) what is the effect of roasting type of coffee, and 5) what is the effect of the duration of brewing tea? All of these scenarios represent typical situations encountered daily in our lives.

## Methods

Generic sample collection method; applied to all of the samples that have been prepared by the methods outlined below. Solid samples (defined here as tomato, yogurt, coffee beans, tea leaves and meat) were collected into one 2 ml round bottom tube (Qiagen) pre-filled with 1.0 ml room temperature ethanol-water (95:5 *v/v*; Ethyl alcohol, pure, 200 proof (Sigma-Aldrich) and Invitrogen UltraPure™ Distilled Water) (for extraction) and one empty 2 ml round bottom tube (as an archive). All solid sample tubes were weighed before and after sample collection and the final sample weight was recorded in the metadata, unless otherwise noted. Samples were fully submerged in ethanol prior to being frozen at −80°C. Liquid samples (defined here as brewed coffee, brewed tea, and milk) were collected into 2 identical empty 2 ml round bottom tubes (Qiagen). After sample collection all samples were frozen and stored at −80°C until downstream sample preparation for mass spectrometry based metabolomics. Solid and liquid samples for each sample type were sampled according to the following schema:

Meat samples (G1): Refrigerated, ground turkey and beef were purchased from grocery stores Trader Joe’s and Ralphs. Three packages of organic products for the beef and 3 packages with the labeling “without antibiotics and growth hormones” for the turkey as well as three packages of products without any “organic” labeling for both meat types were selected. Each package of product was sampled into two petri dishes: one of them was spiked with tetracycline (15 mg/mL tetracycline solution in 70% EtOH (Teknova, Hollister, CA) was diluted with distilled water) to make the final concentration of residual tetracycline to be 300 ppb, while the other was treated with the vehicle (70% EtOH). At 0, 24, 48, 72, and 96 h after treatment, a sample from each meat was collected using ethanol-sterilized forceps and then placed in either a) a tube for archival storage or b) a tube for mass spectrometric analysis.

Tomato samples (G2): Three biological replicates of tomatoes were sampled from distinct sources: organic cherry tomatoes from Vons, non-organic cherry tomatoes from Vons, organic cherry tomatoes from Trader Joe’s, tomatoes from the UC San Diego Farmer’s Market, organic cherry tomatoes from Whole Foods, and cherry tomatoes from a home garden in Los Angeles County, as well as sun-dried tomatoes from Whole Foods and canned tomatoes from Whole Foods. Tomatoes were stored at 4 °C for 20 hours prior to sample collection. For the ripening study, organic tomatoes from Vons were allowed to remain at room temperature with moderate amount of sunlight and absence of ethylene source for 0, 1, 2, 3, 4, and 5 days in biological triplicate. The ripening was arrested by placing the fruit in 4 °C within double Ziploc bag enclosure to avoid acquisition of external compounds. Three technical replicates of each tomato sample were collected using ethanol-cleaned scalpel blades. Samples approximately 3×3×3 mm by size were collected such that each included an equivalent proportion of peel and parenchyma but excluding seeds and placed in either a) a tube for archival storage or b) submerged in ethanol in a tube for mass spectrometric analysis.

Coffee (G3): Coffee was purchased from different roasteries (Perro Negro Coffee Roasters, Eich, Germany; Bird Rock Coffee Roasters, San Diego, USA; Bristol Farms, San Diego USA; Ralph’s, San Diego, USA; Whole Foods, San Diego, USA; Trader Joe’s, San Diego, USA; World Market, San Diego, USA; Illy, Italy; Moniker Coffee Co., San Diego, USA; and Von’s, San Diego, USA) in either ground or whole bean form. Whole beans were ground with a commercial coffee grinder (Hamilton beach fresh-grind #80335). 15 g coffee powder was then brewed with a “french press” brewing device (AeroPress) using 30 mL purified drinking water (Nestle Pure Life) at around 94 °C. After incubating the coffee powder and water for approximately 2 minutes inside the brewing device, coffee was pressed through disposable micro-filters (AeroPress) by hand with moderate pressure. Approximately 1 mL of brewed coffee was poured into the duplicate 2.0 mL tubes and frozen at −80 °C. For solid samples of ground coffee, approx. 0.5 g of coffee was transferred into the sample tubes for archival and extraction for comparison. Two biological replicates for each sample were obtained.

Milk/yogurt samples (G4): Pasteurized whole milk (Horizon Organic Vitamin D Milk) and three different brands of yogurt (Oikos Plain Greek Nonfat Yogurt, Voskos Plain Greek Yogurt, and Kroger Plain Nonfat Greek Yogurt) were chosen. Fermentation was initiated by heating whole milk to 82 °C, followed by cooling to 42 °C. Exactly 25 mL of the milk was then inoculated with 1 mL of each of the store bought yogurts, in sterile 50 mL conical tubes to yield three experimental fermentation conditions. The original milk and the three yogurts were used as controls against the fermented samples, for a total of seven conditions, with three biological replicates for each condition. Samples were sterilely collected by transferring 500 μL of liquid into empty 2.0 mL tubes (Qiagen) and 500 μL of liquid into another 2.0 mL tube (Qiagen) already containing 1 mL of 95% ethanol. The control milk group, that did not have yogurt added, was sterilely sampled at the same time points by transferring 500 μL of liquid each into two empty 2.0 mL tubes. All 21 experimental conical tubes were then covered with aluminum foil and left to incubate at room temperature. Samples were collected starting from the initial inoculation approx. every 12 hours thereafter through 58 hours total (0 hours, 11 hours, 24 hours, 35 hours, 47 hours, and 58 hours), as described above.

Tea samples (G5): Twelve teas (Numi rose white, Allegro tropical white, Prince of Peace oolong, Allegro oolong, Charleston Tea Plantation green, Higgins & Burke green, Lipton matcha, Salada matcha, Charleston Tea Plantation black, Higgins & Burke black, Numi pu’er and Allegro pu’er) representing six tea varieties (white, oolong, green, matcha green, black, and pu’er) were sampled. Tea samples were prepared in two ways. Firstly, tea leaves were removed from the tea bags, divided into triplicates, weighed and placed into a 2.0 mL extraction tube containing room temperature ethanol-water (95:5 *v/v*) and an empty tube for sample archival. The second way in which tea samples were prepared emulates the brewing process. Tea samples were extracted, *i.e.* brewed, in their respective tea bags in triplicates using 200 mL of 95°C bottled water (Nestle) in disposable paper cups (Chinet Comfort Cups). The water was heated with an electric kettle. Aliquots of tea (1000 μL) undergoing the brewing process were taken at 0.5 min, 1 min, 4 min, and 4 hrs after the addition of 95°C water. Aliquots were placed into 2.0 mL tubes and then placed on dry ice. The temperature of the water was measured throughout the extraction, *i.e.* brewing (mean °C of duplicate measures ± reported accuracy of thermometer of 0.1 °C): 0.5 min, 81.5°C; 1.0 min, 81.0°C; 4.0 min, 73.5°C; and 240.0 min, 23.0°C. Extraction blanks (room temperature ethanol-water and 95°C water) were collected in duplicate as controls while monitoring water temperature (1000 jL aliquots collected in 2.0 mL tubes).

### General sample preparation methods for metabolomics

All five sample types were processed with the methods described below. Sample processing [extraction, centrifugation and resuspension]: Solid samples in 1.0 ml of 95% EtOH were thawed over wet ice for 30 minutes. A sterile stainless steel bead was added to each sample and samples were homogenized for 5 minutes at 25 Hz on a tissue homogenizer (QIAGEN TissueLyzer II, Hilden, Germany). Homogenized samples were swabbed with a sterile barcoded cotton swab (BD SWUBE™ Applicator) for future analyses. 100 μL of prechilled extraction solvent (100% EtOH, Sigma-Aldrich) was then added to each well. Liquid samples were thawed over wet ice, gently mixed by inversion and 100 μL were pipetted into a prefilled 96 well deep well plate containing 900 μL prechilled 100% EtOH.

All solid samples were incubated at −20°C for 40 minutes and liquid samples at −20°C for 20 minutes and centrifuged (Eppendorf centrifuge 5418, Hamburg, Germany) for 15 minutes at maximum speed. 400 μL of ethanol extract was carefully transferred to a 96 deep well plate and dried down overnight by centrifugal evaporation (Labconco Acid-Resistant Centrivap Concentrator, Missouri, USA). Prior to LC-MS/MS analysis, samples were resuspended in 50% methanol with 2 μM sulfadimethoxine, as resuspension standard. Samples were vortexed for 2 minutes, followed by 5 minutes sonication (Branson 5510, Connecticut, USA) in a water bath, before centrifuging (Thermo SORVALL LEGEND RT, Germany) for 15 minutes at 4°C. Samples were transferred into a 96 well shallow well plate, and run at 1x dilution or diluted 5x prior to analysis. Samples were centrifuged for 5 minutes before placing on the autosampler and 5 μL of extract were injected for each sample.

### Sample metadata curation

Metadata were entered manually for 704 samples of the above described food and beverages. Images were used to capture key sample information including unique barcode IDs, packaging information and time of sample collection. Metadata consists of 142 different descriptive categories including, but not limited to, information about the following: ingredients, packaging type, location of food production, location of sample collection, store and brand names, UPC codes, NDB numbers and descriptions, cheese and dairy types, fermented and non-fermented foods, botanical definitions and genus names of plant samples, conventional vs organically produced, type of animal meat, and presence of common allergens and additives. Sample information entries were standardized by use of a metadata dictionary that explained the type of information needed for each category as well as the correct formatting.

### Mass spectrometry data collection

Food sample extracts were chromatographically separated with a UltiMate 3000 UPLC system (Thermo Scientific, Waltham, Ma). Reverse phase chromatographic separation was achieved with a Kinetex C18 column (100×2.1 mm) packed with 1.7 μm particles from Phenomenex (Torrance, CA, USA), fitted with a guard cartridge. The column compartment was held at 40°C, with a constant flow rate of 0.5 mL/min. A linear gradient was applied: 0-1.5 min isocratic at 5% B, 1.5-9.5 min 100% B, 9.5-12 min isocratic at 100% B, 12-12.5 min 5% B, 12.5-14 min 5% B; where mobile phase A is water with 0.1% formic acid (v/v) and phase B is acetonitrile 0.1% formic acid (v/v) (LC-MS grade solvents, Fisher Chemical). The UPLC system was coupled with a Maxis Q-TOF mass spectrometer (Bruker Daltonics, Bremen, Germany) controlled by Otof Control and Hystar software packages (Bruker Daltonics, Bremen, Germany) and equipped with an ESI source; MS spectra were acquired in positive ionization mode with mass range 50-1500 *m/z*. The instrument was externally calibrated to 1.0 ppm mass accuracy with ESI-L Low Concentration Tuning Mix (Agilent Technologies, Waldronn, Germany) twice daily. During the run, hexakis (1H, 1H, 2H-difluoroethoxy)phosphazene (Synquest Laboratories, Alachua, FL) with *m/z* 622.029509, was used for lock mass correction. For data dependent acquisition the five most abundant ions per MS1 scan were fragmented and the spectra collected. MS/MS active exclusion was set after 2 spectra and released after 30 seconds. A fragmentation exclusion list was set: *m/z* 144.49-145.49; 621.00-624.10; 643.80-646.00; 659.78-662.00; 921.0-925.00; 943.80-946.00; 959.80-962.00 to exclude known contaminants and infused lockmass compounds. A process blank was run every 12 samples; a standard mix [Sulfamethazine 10 mg/l, Sulfamethizole 10 mg/l, Sulfachloropyridazine 10 mg/l, Sulfadimethoxine 10 mg/l, Coumarin-314 20 mg/l] was injected 1-2 times per plate of samples [a 96 well plate contained 8 blanks and 88 samples max]. These intensity and RSD of the standards are plotted in **Supplementary Figure 12**.

### QC assessment of the data collection

Bruker raw data files were lock mass corrected (*m/z* 622.0290) and converted to .mzXML files using Bruker DataAnalysis software. Raw data files were manually inspected in Bruker DataAnalysis software. Files that did not contain the resuspension standard (sulfadimethoxine, [M+H]^+^, *m/z* 311.0809) were not included for further analysis. The turkey and beef samples had significant lipid carryover, which was minimized by running two wash cycles and one blank in between each of the meat samples. Caffeine was observed as a contaminant between runs due to the high intensity in some samples, and this carryover was removed during feature finding, described below. Files were organized into samples, wash, controls, blanks and the data were uploaded via GNPS and stored in MassIVE (https://massive.ucsd.edu/): Meat (G1): MSV000082423; Tomato (G2): MSV000082391; Coffee (G3): MSV000082386; Milk/yogurt (G4): MSV000082387; Tea (G5): MSV000082388.

### Molecular networking and small molecule annotations

For molecular networking parameters were set to a minimum requirement of 4 ions to match and a cosine score of >0.7. Parent mass tolerance was 0.1 Da and MS/MS was set to 0.1 Da (these parameters were used as many reference type spectra are low resolution). The library search was performed with min match peaks of 4 and a cosine >0.7. Due to the different small molecule compositions for each food, the annotations of all individual food analyses are impacted differently as recently shown with Passatutto, a false discovery rate (FDR) estimator (Scheubert et al., 2017). Passatutto was used to estimate FDR for the annotations with our settings for each of the five sub-analyses. Passatutto uses a decoy database created using fragmentation trees and rebranching of fragments to estimate the FDR. With these analysis parameters the estimated FDR of annotations based on spectral matching, at level 3, for the meat data is at 1.5%, for the tomato data 4.8%, coffee 0.09%, milk to yogurt 0.5%, and 0.2% for tea.

### Feature finding

MS1 feature detection was performed using MZmine 2.32 in batch mode. Only features linked to MS/MS were kept in the final output. For this purpose mzXML files were imported and cropped based on retention time (0-11.5min). The mass detection noise level for MS1 was set at 1.0E3 and 5.0E2 for MS/MS. Chromatograms were built with a min time span of 0.01 min, a min height of 3.0E3 and a *m/z* tolerance of 20.0 ppm. Chromatograms were deconvoluted using local minimum search (chromatographic threshold: 20%; search minimum in RT range (min): 0.05; minimum relative height: 25%; min absolute height: 2.0E3, min ratio of peak top/edge: 1 and peak duration range (min) of 0.05 − 3.00). The *m/z* range for MS/MS scan pairing was 0.05 Da and the RT range for MS/MS scan pairing was 0.1 min. Isotopic peaks were grouped with an *m/z* tolerance of 20.0 ppm, a retention time tolerance of 0.1 and a maximum charge of 4. Features were then aligned with a *m/z* tolerance of 20.0 ppm, weight or *m/z* 75% and weight for RT 25%, with a retention time tolerance of 0.4 min. Gap filling was then performed with an intensity tolerance of 10.0% and a retention time tolerance of 0.3 min. Feature tables were then filtered to include only features that contain a minimum of 2 peaks in a row and which have MS2 scans in at least one sample.

The filtered feature tables were then further processed to remove MS1 features (within 10 ppm mass error) associated with the lock mass ([M+H]^+^, *m/z* 622.0290 and [M+Na]^+^, *m/z* 644.0109), the resuspension standard (sulfadimethoxine, [M+H]^+^, *m/z* 311.0809), the standard mix, described above, as well as carry-over contamination from caffeine ([M+H]^+^, *m/z* 195.0877). Feature tables were concatenated with metadata, based on the MS filename and used for MS1 analysis. The final feature tables used for PCoA analysis and heatmaps were uploaded to GNPS (http://gnps.ucsd.edu) to create each respective MassIVE accession link for public access.

### PCoA

We used Principal coordinates analysis (PCoA) to observe broad molecular patterns and trends within the data PCoA takes a dissimilarity matrix as input and aims to produce a lowdimensional graphical representation of the data such that samples closer together have smaller dissimilarity values than those further apart. PCoA plots consist of orthogonal axes where each axis (PC1, PC2, PC3) captures a percentage of the total variance. For PCoA, the signal intensities of the features were normalized with Probabilistic Quotient Normalization (PQN) (Ejigu et al., 2013). The PCoAs were calculated using the Canberra dissimilarity metric using QIIME (Caporaso et al., 2010) and visualized in EMPeror (Vázquez-Baeza, Pirrung, Gonzalez, & Knight, 2013).

### Heatmaps

Heatmaps were created from the filtered and preprocessed feature tables, comprising both overall features as well as only features with a GNPS library hit. Jupyter notebooks used to create the heatmaps are publically available at: https://github.com/DorresteinLaboratory/supplementary-MolecularChangesInFood

## Results

A wide variety of foods including meat, tea, coffee, tomato and home fermented yogurt were sampled based on processing types and/or longitudinal changes to determine molecular variation associated with each processing method. **Figure 1** highlights some representative examples of images associated with the specific foods that were sampled. The numbers on the tubes indicate the barcode number associated with each file, which was used to track the information and metadata for the entire project. **Figure 1a** shows an example of ground beef at 3 days left at room temperature (a part of a 5 day time course to investigate meat spoilage). The discoloration of the meat is non-uniform. The next sample type is tea, where twelve teas were subjected to brewing for 0.5 min, 1 min, 4 min and 240 min. A representative sample point of one tea at 1 min is shown in **Figure 1b**. The third sample type is tomatoes (**Figure 1c**). Both the tomato origin and impact of time of storage at room temperature on the molecular make-up were investigated. The fourth sample type studied was the home fermentation of yogurt, over 6 days, including controls of the milk and initial yogurts containing live active cultures. One example for yogurt fermentation is shown in **Figure 1e**. Finally, we assessed different roasts of coffee (the packaging for the medium/dark roast is shown in **Figure 1d**). Each of the samples were subjected to extraction as outlined in the methods and the resulting extracts were subjected to LC-MS/MS-based mass spectrometry. To obtain an overview of the data, we created PCoA plots, heatmaps and molecular networks.

**Figure 1.**
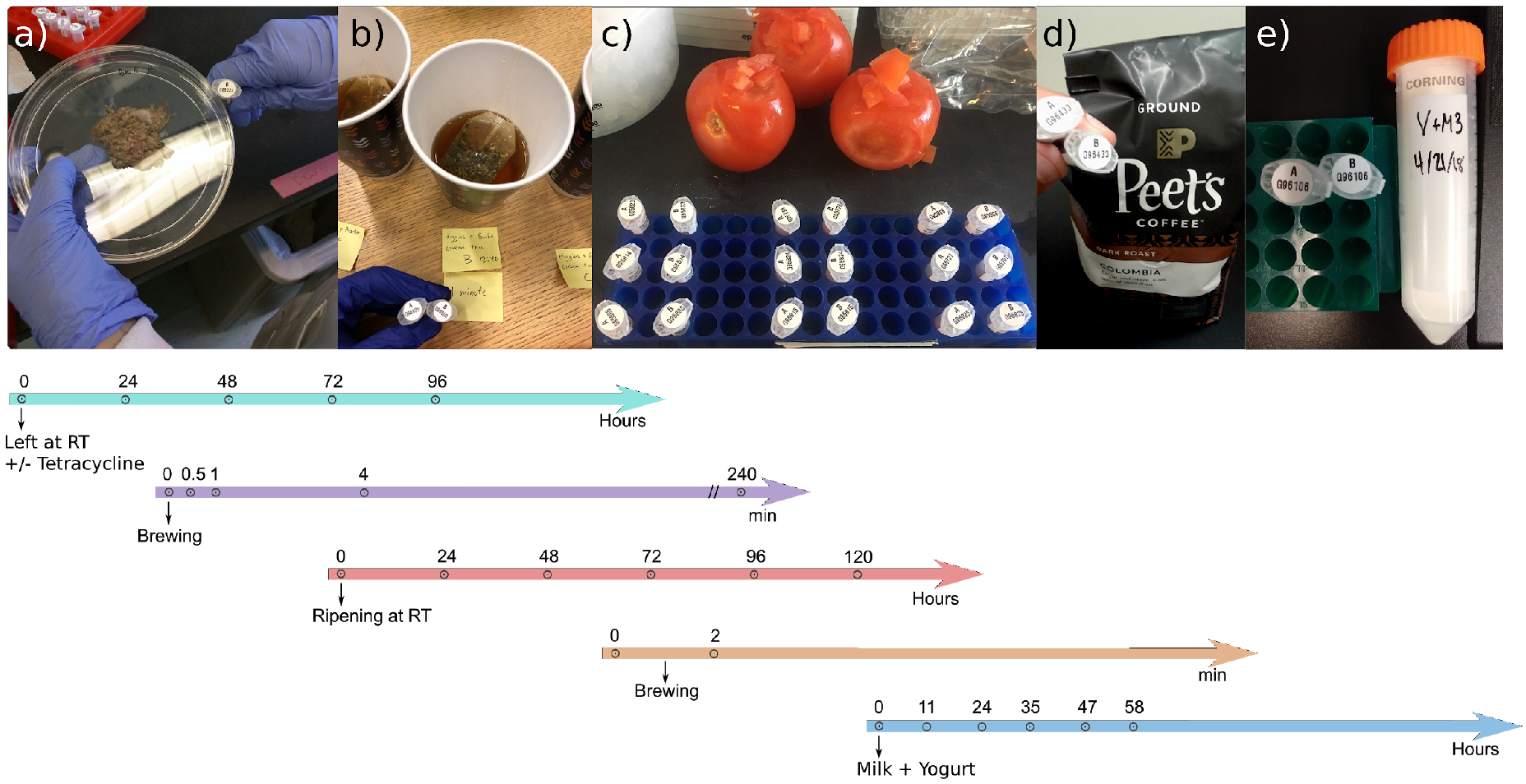
Representative images of the foods sampled. a) Ground beef at 3 days storage, b) Tea brewed at 1 min, c) Tomato slices from a local Farmer’s market, d) Coffee, e) Yogurt preparation from milk. The timelines in the lower panel indicate the sampling times by the numbers indicated. RT denotes room temperature and // denotes a time break.

### PCoA

A global PCoA analysis of all datasets revealed clustering based on the different sample types (**Figure 2 and movie S1**). Tomato and meat samples form tight groups, red and blue respectively, while the tea has two groups representing the solid (tea leaves) and brewed samples. Coffee has three groups due to extractions of brewed samples, direct extraction of ground beans and then extraction of the beans themselves. The dairy samples differentiate between the milk and the samples containing yogurt cultures. To further visualize details within a sample type, PCoA analyses were performed on each dataset independently.

**Figure 2.**
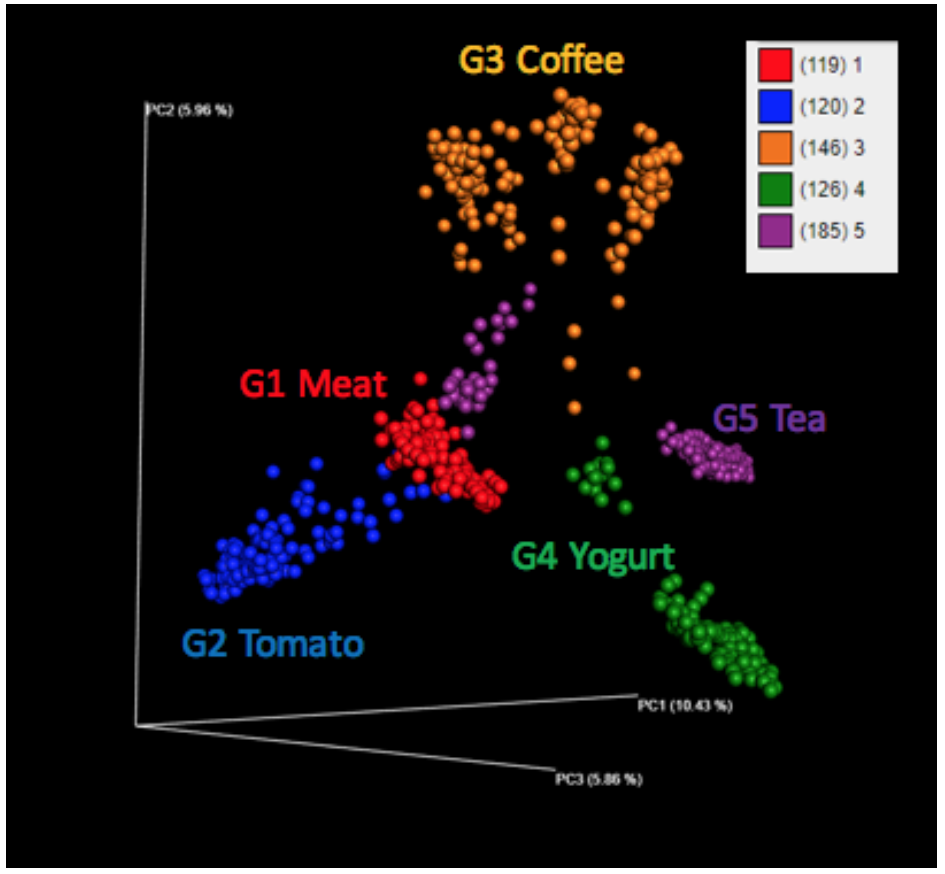
Global PCoA analysis to understand the molecular relationships among all the samples analyzed. PC1 (10.43%); PC2 (5.96%); PC3 (5.86%). As a 2D image, the PCoA plot does not reveal the relationships clearly, a movie rotating this image is provided as supporting information. In parentheses the number of samples for each group are shown.

The PCoA analysis of the yogurt and milk samples shows distinct grouping between yogurt and milk samples (**Figure 3a-3d**). In fact, each brand of yogurt is distinct despite containing similar live active cultures and ingredients (yogurt 1 = Sun Valley Dairy; yogurt 2 = Oikos; yogurt 3 = Kroger). Oikos and Sun Valley Dairy contain the same live active cultures (*S. thermophilus; L. bulgaricus; L. acidophilus; Bifidus; L. casei*) whereas Kroger contains *L. acidophilus, B. bifidum, and L. casei*. The home fermentation time courses of the milk inoculated with the different yogurts as starter culture are displayed in **Figure 3b-3d**. The fermentation process and associated molecular changes can be visualized by all three home ferments becoming more yogurt-like, as evidenced from the later time points becoming closer to yogurt than the starting milk in PCoA space. For example, **Figure 3c** nicely illustrates the transition of the home ferment (milk + starter yogurt) through time, becoming more similar to the original starter culture. Sun Valley Dairy contains Grade A pasteurized milk and cream, in addition to nonfat milk found in the Oikos, possibly contribute to the difference between these yogurts and the corresponding home ferment.

PCoA analysis of ground turkey and ground beef left at room temperature and the impact of tetracycline are shown in **Figure 3e-3g**. In PCoA space the attribute that most differentiates the samples is the type of meat (**Figure 3f**). Samples with and without tetracycline addition change similarly over time, indicating that based on this multivariate analysis, the addition of tetracycline does not greatly impact the aging process (**Figure 3g**). Surprisingly, even after leaving the meats out at room temperature for 5 days and the development of a significant emanating odor from the samples, no trend could be spotted in the PCoA. A greater change may be detected using other methods such as GC-MS that would detect volatile compounds. As **Figure 1a** reveals, the aging process is non-uniform and thus the experimental variation of the samples within the same time points appears to be larger than the overall molecular variation associated with the 5 day aging process.

PCoA analysis of the tomato samples revealed that both source (**Figure 3h**) and storage time (**Figure 3i**) affect the molecular composition of tomatoes. As expected, canned and sundried data occupy very different PCoA space than fresh tomatoes. It is also notable that differences exist for fresh tomatoes, with those from Farmer’s market most closely resembling home garden tomatoes and all store-bought tomatoes resembling one another, whether organic or not. When organic tomatoes are left at room temperature they occupy the bottom left corner in **Figure 3i** and gradually change to the lower right over the course of 6 days suggesting that there are major molecular changes over this time period and upon inspecting **Figure 2** it appears that these changes are larger than the changes in meat over the same period. Notably, despite the magnitude of these molecular changes the tomatoes did not change at all in either their appearance or smell.

PCoA analysis of the coffee revealed a clear trend among the sample type liquid “brewed coffee” or solid “ground coffee” (**Figure 3j-l**). If the coffee sample was a liquid from brewing and the brew was extracted, then the sample appeared on the left side of the PCoA, while extracts of the ground beans (picked up with clean spoon) or the cut beans (with sterile knife) themselves directly appeared on the right. Besides sample type, the data suggest there is clustering based on the roasting type, as there are clusters associated with clustering of dark roast, light roast and the medium roast. This is particularly noticeable when the coffee is extracted from the ground beans and/or directly from the beans (**Figure 3k**).

PCoA analysis of tea samples, **Figure 3m-3o**, revealed unambiguous differentiation of solid from liquid samples, prepared using room temperature ethanol solution and 95°C water, respectively. Note that water samples were most differentiated from the solid extract samples along PC1. Twelve different teas were sampled over time (0.5 min, 1 min, 4 min, and 240 min) to emulate the brewing process (**Figure 3m**). A water-only control at the same time points did not change (**Figure 3m**) and direct extractions of the solid teas are also shown. Interestingly, the samples appeared most similar to water blanks in the earliest time points and became more similar to the solid samples over time (along the PC1 axis which explained 25.9% of total variance), independent of tea type, indicating continued release of compounds from the leaves. The kinetics of tea extraction were similar for all teas ‑ interestingly, the observed chemical differences between 240 min and 4 mins were minor for all teas which supports a steeping time rationale which appears to be sufficiently effective for tea extraction of phytochemicals. A slight deviation in overall kinetic trend was observed for oolong. The first two time points (0.5 and 1 min) appeared to be more similar to the blanks than other teas at the same time point. Differences based on tea type were also observed. White, green, matcha, and black tea liquid samples were more similar to each other than to oolong and pu’er, which were differentiated along PC3 (6.81% of total variance); **Figure 3n** and **3o**, illustrate the clear differences between Chinese teas (oolong and pu’er), **Figure 3n**) and the American and British teas (**Figure 3o**). Although PCoA enables the detection of overall trends, PCoA does not enable looking at changes in levels of the individual molecules.

**Figure 3.**
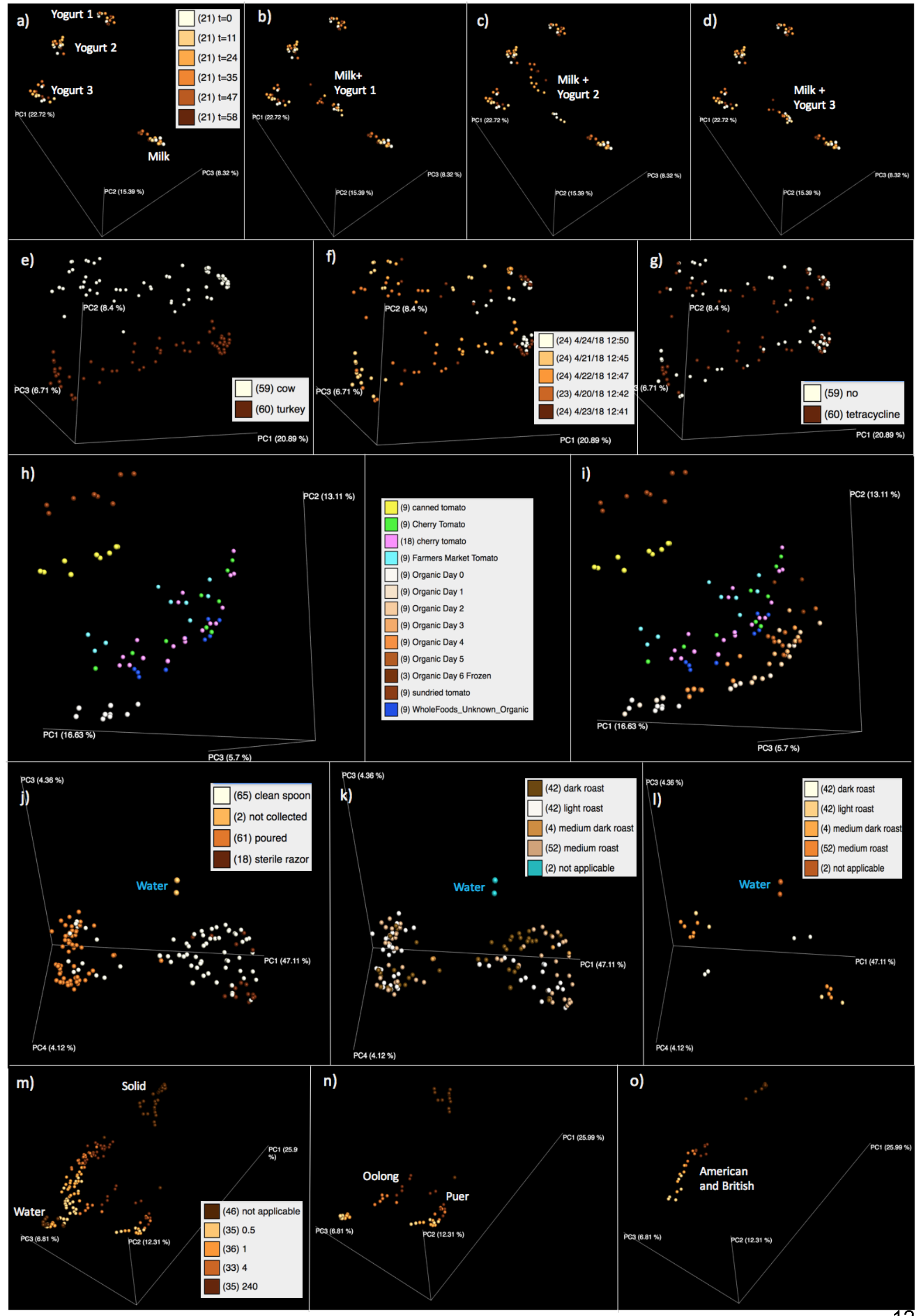
PCoA plots for the individual food types, color coded by metadata categories to visualize key drivers in molecular patterns. The three store bought yogurts containing live active cultures, the milk and the home ferments using the different yogurts as starter culture show distinct groupings. The spheres are colored based on fermentation time from 0 to 58 hrs (a-d). The meat samples separate by animal type (e), duration left at room temperature (f), but do not show a clear trend based on tetracycline addition (g); tomato samples display differences based on source (h) and storage time (i); coffee samples group based on collection device, which tracks with brewed coffee vs. ground beans (j); the impact of the roast type is also depicted in (k) and (l). Tea samples differentiated based on whether they were extracted with ethanol (tea leaves) or first extracted with water, over a range of brewing times (m); different Chinese tea varieties group separately (n) from the British and American teas (o).

### Heatmaps

We created heatmaps to visualize molecular changes driving differences between samples for the time course experiments of tea brewing, yogurt fermentation, tomato ripening and improper meat storage and gain more insight into groups of features that behave similarly over time or in different sample types. In addition to a PCoA, heatmaps provide a visual overview of the data to give more detailed insights behind molecular changes driving the differences between sample types and within sample types. Because the tea and the milk-to-yogurt had the largest changes in abundances of groups of molecules they are shown in **Figure 4**. Other heatmaps are shown in the Supplementary Information (**Supplementary Figure 1**-**4**). Consistent with the PCoA analysis, we observe different metabolite profiles between solid and liquid samples in tea (**Figure 4a**). Furthermore, we observe that relative intensity of molecular features increases with extraction time independent of the tea type. We assessed the correlation of relative intensity per feature and tea type with extraction time. In tea this resulted in a total of 2,045 significantly correlated features (*spearman correlation, p-value* < 0.05). **Figure 6c** highlights selected molecular features for which we obtained a putative structure annotation through GNPS library matching. For example, we observe that the relative intensity of procyanidin B and theaflavin increase over time (*Kruskal-Wallis, N=6, p-value* ranging from 0.01 to 0.02, *between brewing times 0.5 and 240*). We also assessed the correlation of relative intensity per feature and home ferment with different yogurt inoculums over time. For the Kroger yogurt, this resulted in a total of 1,587 significantly correlated features (*spearman correlation, p-value* < 0.05). **Figure 5b** highlights selected molecular features for which we obtained a putative structure annotation through GNPS library matching. For example, we observe that the relative intensity of 4-O-beta-galactopyranosyl-D-mannopyranose decreases over time for each yogurt type individually as well as overall (*Kruskal-Wallis, N=9, p-value=0.0023, between 0 and 58 hrs*).

Molecular changes during meat (beef and turkey) storage over five days were also visualized (**Supplementary Figure 2a**). When comparing antibiotic vs non antibiotic treated meat (both beef and turkey), the overall molecular differences as seen in PCoA space do not vary much. However, there are some specific low intensity molecules that change, although with minimal differences due to the addition of tetracycline, consistent with the observations from the PCoA. We do observe differences between organic and non-organic beef. For example, in the non-organic beef, oleoyl-taurine increases during the 5 days and does not appear by day 5 in the organic samples, while the levels of acetyl-carnitine decrease in the non-organic beef but are consistent across all time points for the organic beef. In the turkey the rate of appearance of oleoyl-taurine and rate of disappearance of acetyl-carnitine are only slightly different (**Supplementary Figure 2b**). The spectral match with parent mass difference 0.000 Da and very strong cosine match of 0.84, to the fungal molecule, termitomycamide E (Choi et al., 2010), increases over time, the presence of three analogues with mass differences pointing to different acyl chain lengths, and minor suppression by tetracycline appears to have been detected and would be consistent with increased microbial loads (**Supplementary Figure 4**).

Molecular differences between tomato samples were most striking when comparing sun dried, canned and fresh tomatoes. In the heatmap visualizing molecular changes during the ripening of fresh tomatoes (**Supplementary Figure 1**) no clear-cut large scale patterns were observed. During the ripening process some individual molecular features were found to decrease in their relative abundance. For example, 5’-methylthioadenosine, a molecule, which can be used to produce ethylene, a key ripening hormone for plants (North, Miller, Wildenthal, Young, & Tabita, 2017) was found to decrease significantly in its relative abundance over the 5 day time course. Also plant flavonoids (including level 3 annotation of naringenin) and tomatidine, a tomato-specific alkaloid, were found to decrease significantly in their relative intensity over time. This is informative as many of the healthy properties assigned to polyphenol-containing food are attributed to molecules like naringenin and it indicates that the nutritional value of tomatoes may change over the time period we typically store tomato fruits at home.

**Figure 4.**
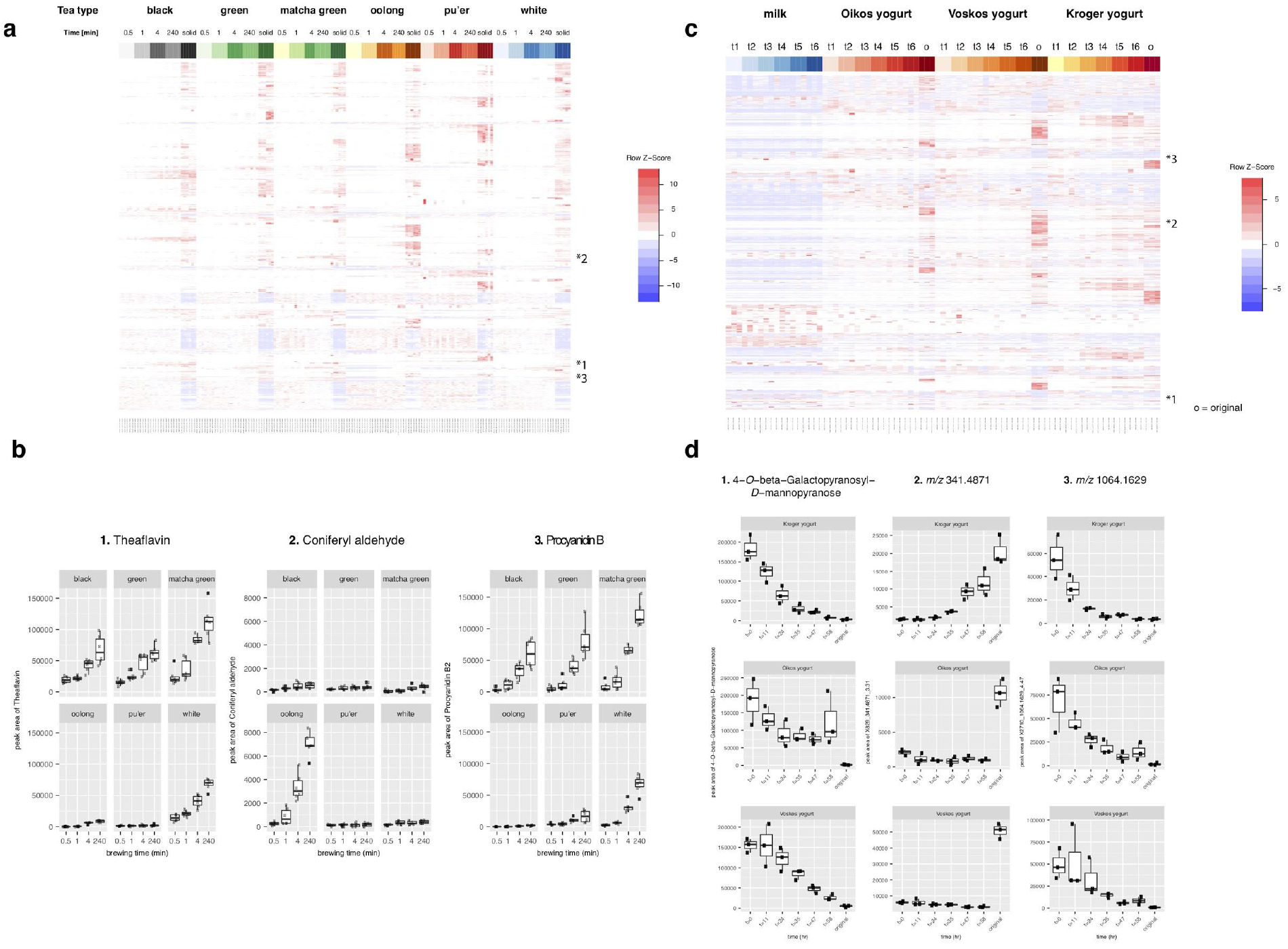
Metabolites changing over tea extraction time and during the fermentation process from milk to yogurt. a) Heatmap showing tea metabolites changing over extraction time across different tea types. b) Specific metabolites increasing significantly in their relative intensity during tea extraction time. c) Heatmap showing metabolites changing during the fermentation process from milk to yogurt across different yogurt brands used as inocula, as well as the milk as control. d) Metabolites increasing or decreasing significantly during the fermentation process across different home ferments. Metabolite annotation was performed through mass spectral molecular networking and spectral matching to reference spectra as indicated below.

### Molecular networking and annotations

To further explore specific molecules and molecular changes within each food type, we subjected all LC-MS/MS data to mass spectral molecular networking. Mass spectrometry of the tomato samples (120) resulted in 71,430 MS/MS spectra, 62,263 passed the filtering for a minimum of 4 ions and a minimum of two identical MS/MS spectra in the data set and this condensed to 2,611 unique spectra that are presented as nodes (**Supplementary Figure 5**). 212 of the nodes had an annotation. This is an 8.1% annotation rate and with an FDR for spectral matches estimated using Passatuto to be 4.8% (Scheubert et al., 2017). All annotations are level 2 or 3 according to the 2007 metabolomics standards initiative (Sumner et al., 2007). For the milk to yogurt analysis, the 126 samples resulted in 78,203 MS/MS spectra, 63,241 passed the minimal requirement of four ions and minimum of two identical spectra (**Supplementary Figure 6**). Post clustering identical spectra, 4,142 nodes remain. 147 of the nodes had spectral matches against the libraries (3.5% annotation rate, FDR 0.5%). The coffee analysis included a total of 146 samples that resulted in a total of 50,929 MS/MS spectra. After filtering, 42,752 MS/MS spectra remained that condensed to 1,460 unique spectra in **Supplementary Figure 7**. Of the 1,460 unique spectra, 72 had spectral matches to the reference libraries within a cosine of 0.7. This is a 4.9% annotation rate and with an FDR estimated to be 0.09%. The meat analysis included 119 samples, resulting in 72,083 MS/MS spectra, 54,663 of which passed the filtering step (**Supplementary Figure 8**-**9**). Merging all identical spectra resulted in 5,035 unique spectra of which 313 were annotated (6.2% annotation rate, FDR 1.5%). Finally, the tea analysis had 185 samples resulting in 50,547 MS/MS spectra, 44,505 of which passed the filtering (**Supplementary Figure 10**-**11**). After merging identical spectra, 1,834 unique MS/MS spectra comprised the molecular network with 207 annotations (11.2% annotation rate, FDR 0.2%).

MS/MS belonging to the internal standard sulfadimethoxine was observed in all analyses and correctly annotated through GNPS library matching. Molecules annotated as the amino acids tryptophan and phenylalanine as well as phospholipids were widely distributed across all samples and different food types. Other putatively annotated molecules were found to be food-specific.

In the tomato samples (**Figure 5a**), many spectral matches to chlorogenic acid derivatives and flavonoids were detected, both compounds indeed commonly found in different tomato cultivars (van der Hooft, Vervoort, Bino, & de Vos, 2012, Floros et al., 2017). Moreover, a molecular family of tomatidine-related molecules was observed throughout all tomato samples. A molecular family is a set of MS/MS spectra that are similar from which the structural relatedness is inferred (Nguyen et al., 2013). As the name suggests, this is a tomato-specific alkaloid that is the basis for glycoalkaloids like tomatine found abundantly in tomato plant leaves and stem and at minor concentrations in the fruits. Similarly, phenylethyl pyranosides are found in all tomatoes, irrespective of processing. 5’-methylthioadenosine was detected in all tomatoes except sundried tomatoes and can be used to produce ethylene, a key ripening hormone for plants (North, Miller, Wildenthal, Young, & Tabita, 2017). The relative concentration of 5’-methylthioadenosine is observed to decrease over the time course of ripening (**Supplementary Figure 1**), while other molecules increase. Only in the sun dried tomatoes did we observe a spectral match to glucose, perhaps added as a sweetener. In both sun dried and fresh tomatoes we detected azoxystrobin, a fungicide used as protectant against fungal diseases in agriculture.

In milk and yogurt, matches to six carbon sugars, disaccharides and oligosaccharides, vitamins and acylated carnitines were observed (**Figure 5b**). In addition, large lipid molecular families, such as sphingolipids, and glycerol conjugated with fatty acids such as monoolein and linoleoylglycerol were annotated. Delvocid, also known as the clinically used antimicotic natamycin, which is a known additive used to preserve dairy products from fungal growth, was detected (Branen, Davidson, Salminen, & Thorngate, 2001), and did not change in relative abundance over time. These annotations are all consistent with the animal, milk and yogurt origin of the samples. However, we also obtained unexpected annotations. The bile acids glycocholic acid and cholic acid formed an annotated molecular family. These were not expected to be observed as they are primarily associated with the gut. Although level 3 annotations, manual inspection of the ions and retention time analysis reveal the data are indeed consistent with these bile acids.

In coffee (**Figure 5c**) we observed caffeine as well as methyl-caffeine and a related compound with a delta mass of *m/z* 14.01 (CH_2_), corresponding to theobromine. Furthermore, we detected several flavonoids and a large number of hydroxycinnamic acids and chlorogenic acids, which are commonly observed in plants (Islam MT, et al., 2018, Clifford, Jaganath, Ludwig, & Crozier, 2017, Pastoriza, Mesías, Cabrera, & Rufián-Henares, 2017, Karpinska, Świstocka, & Lewandowski, 2017, Tajik, Tajik, Mack, & Enck, 2017, Naveed et al., 2018a, Naveed et al., 2018b). In addition, library matching revealed presence of mascarosides, molecules commonly observed upon roasting of coffee (Shu et al., 2014). The ascarosides were noted in the molecular network by *m/z* 162.053, 15.996 and 18.011 gains and losses, corresponding to mass shifts associated with six carbon sugars, oxygen, and water, respectively.

In the meat samples (**Figure 6a**), we observed MS/MS matches to tetracycline displayed as a single node (no related spectra were detected), which were more abundant in the turkey samples. Although tetracycline is commonly used as a growth promoter, here it was added to see the effect of this antibiotic on a 5 day food spoilage test (Granados-Chinchilla & Rodríguez, 2017). We also have spectral matches to carnosine as well as a large cluster of acyl carnitines with five spectral matches to different acylations. The acyl carnitines are predominantly observed in beef when comparing to turkey. We also found a family of N-acyltaurines (NATs), a recently discovered class of lipids (Turman, Kingsley, Rouzer, Cravatt, & Marnett, 2008). **Supplementary Figure 2** shows how after two days storage at room temperature levels of NATs increase whereas levels of acylcarnitines (markers for beta-oxidation) drop, suggesting a change in metabolism over time. Taurine is an organic compound that is widely distributed in animal tissues. Ceramides, component lipids of one of the major bilayer eukaryotic cell membranes, are detected in both beef and turkey but only in beef do they fall below the detection level after 5 days. Their presence marks disintegration of cells within the meat and the lability of ceramides explains their disappearance over time. A spectral match to carnosol, a metabolite from rosemary plants, and connected MS/MS spectra were observed in turkey, but not in beef. Only the packaging of the turkey grown without antibiotics and growth hormones states that rosemary was used, yet it is observed in the conventionally grown as well as antibiotic-free meat. Both dipeptides and *N*-methyl histidine are also detected during the 5 day aging process of the meats. Thus, in general the majority of annotations are consistent with what we would expect to observe in these sample types and we observed changes in molecular compositions emerging after 2 days of storage but only for a small number of molecules.

A large range of phytochemicals were annotated in the tea samples (**Figure 6c** and **Supplementary Figure 10**) including large molecular families associated with flavonoids with spectral matches to puerins, catechins, and apigenin (assignment are putative as isomers are difficult to differentiate in accordance with level 3 metabolite identification (Cuyckens & Claeys, 2004, Sumner et al., 2007, Borges et al., 2018). Mass spectrometry work on tea is primarily done in negative ionization mode; here, using positive ionization mode, we corroborate earlier work by finding molecular families containing the flavonoid aglycones with MS/MS matches to quercetin, kaempferol, myricetin, and (epi)catechin ‑ glycosides that are abundant in tea (van der Hooft et al., 2012) and a large molecular family consisting of glycoside derivatives that have spectral matches to quercetin and kaempferol that are bundled together with chlorogenic acids. Note that the majority of nodes for this family were annotated with GNPS community contributed library hits, indicating that for some compound classes library coverage is increasing due to the growing publicly available spectra. As with coffee, caffeine was annotated in the tea samples as well. Theaflavin, a polyphenol formed during fungal oxidation and its analogues often associated with black tea (Zhang et al., 2018), were detected in white, green, black and oolong tea samples, and as seen in **Figure 4** and **Supplementary Figure 11**, it increases in relative concentration as the tea sits. These annotations are consistent with the known processes that use the polyphenol building blocks to create larger scaffolds like theaflavin, giving black tea its typical brown color. Furthermore, fuzhuanins, polyphenol-derived molecules (Luo et al., 2013) which are beta-ring fission lactones of flavan-3-ols like epicatechin, were found at high abundance in tea.

**Figure 5.**
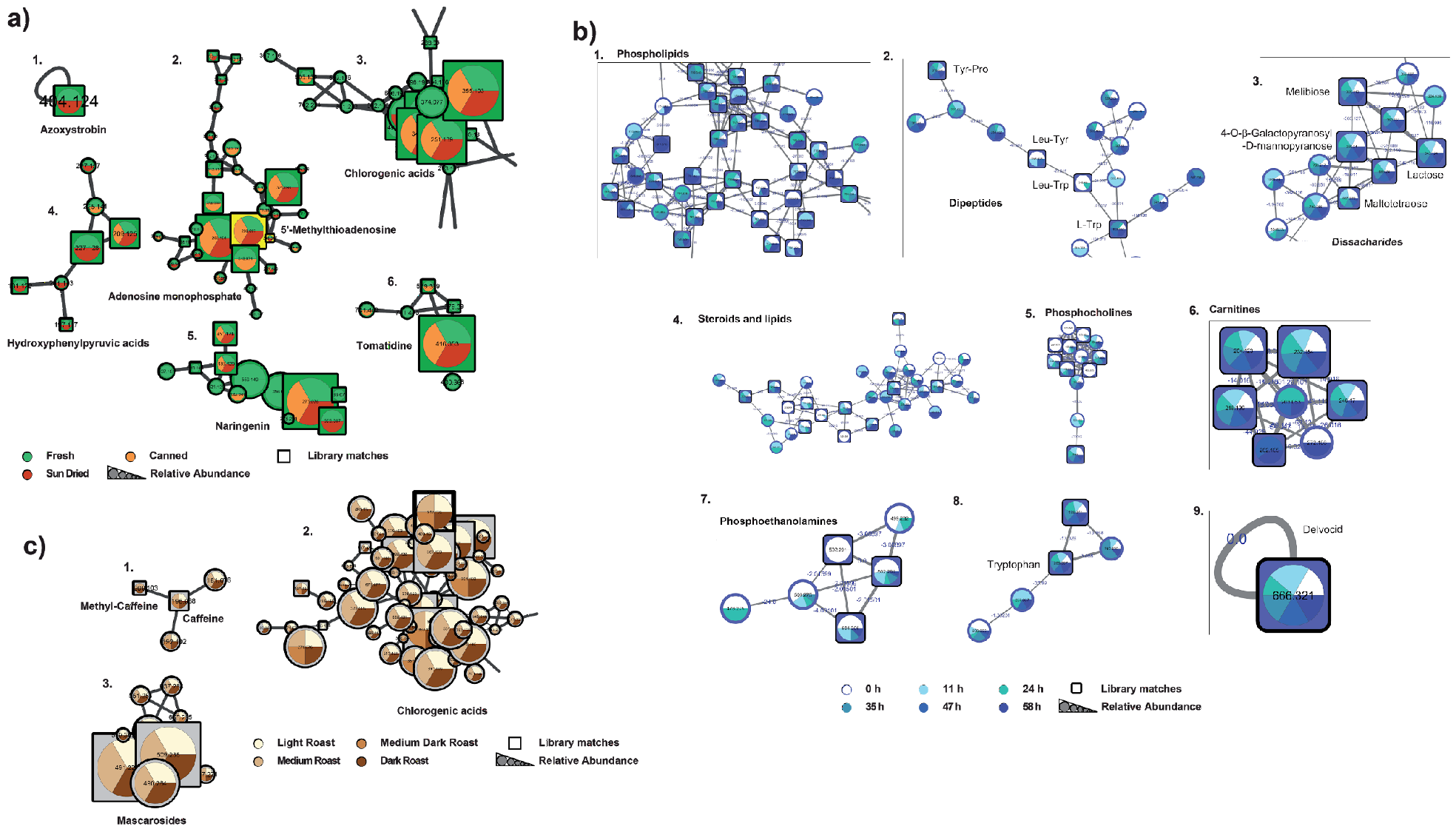
Molecular network clusters of the a) tomato color coded by processing method, b) milk to yogurt, c) coffee data. The clusters are enlarged regions of specific molecular families observed within the full molecular network. The color coding for different samples groups are explained in the figure legend. Node sizes indicated relative precursor abundance and selected library identifications are annotated in the figure and shown through squared node shape. The full size images of the entire network where one can zoom in to the molecular networks can be found as supporting information (**Supplementary Figure 5**-**7**) and the GNPS links to the analysis jobs are provided in the data availability section. All annotations shown are level 2 or 3 according to the 2007 metabolomics standards guidelines (Sumner et al., 2007).

**Figure 6.**
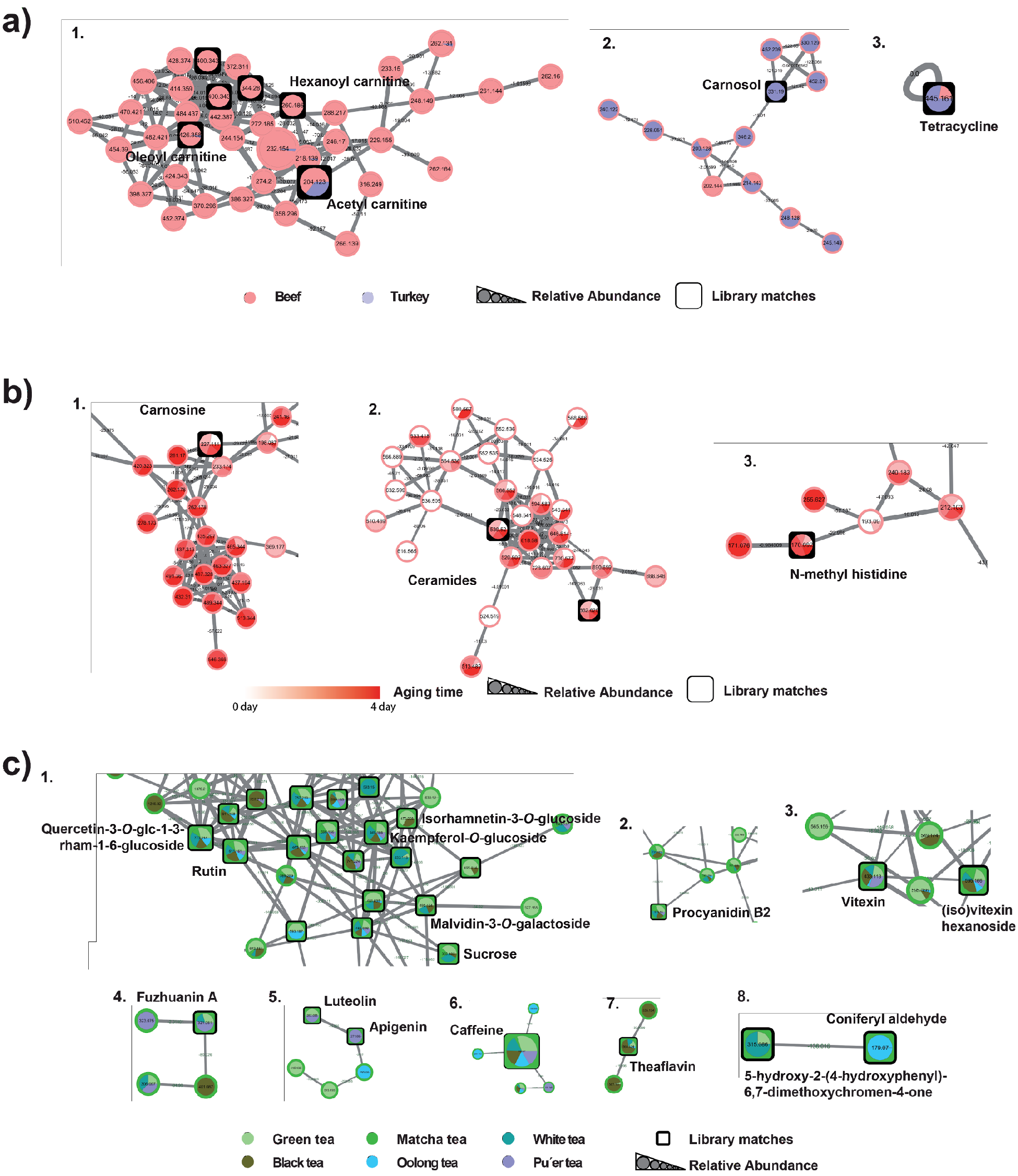
Molecular networks of the data. a) reflect the meat samples color coded by turkey or beef. b) same network as a) but color coded by aging time. c) molecular networks color coded by tea. The insets are enlarged regions of specific molecular families observed within the molecular networks. The full size images of the entire molecular networks where one can zoom in molecular networks can be found as supporting information (**Supplementary Figures 8**-**10**) and the GNPS links to the analysis jobs are provided in the data availability section. All annotations shown are level 2 or 3 according to the 2007 metabolomics standards guidelines (Sumner et al., 2007).

## Discussion

The untargeted mass spectrometry approach coupled with molecular networking allowed us to assess large scale differences between sample type, find molecule-molecule links within and between sample types, and identify different compound classes found within a sample type - all useful for biochemical interpretations. We determined that different foods undergo different molecular changes over time, exemplified by the tea and yogurt time courses. Furthermore, mass spectral molecular networking could identify key metabolites, which differed based on processing type, such as fermentation time in the yogurt samples and brewing time for tea. The heatmaps as well as the molecular networks, while very different visualization techniques, confirm and support each other. For example, theaflavin increases significantly in relative abundance over time which can be visualized in the heatmap (**Figure 4a,b**) as well as the molecular network displaying brewing time (**Supplementary Figure 11**). In **Figure 6c** one can see that theaflavin is connected to two unannotated compounds, which allows us to understand more about the compound family without understanding the exact identity. Furthermore, foods within a group were found to undergo differential molecular changes over time, exemplified by the extraction kinetics of oolong tea deviating from the rest of the tea samples. Two potential explanations for the observed changes in extraction kinetics for tea are hypothesized. The oolong tea bags might affect extraction kinetics; however, the observed differences were observed in both manufacturers’ brands. The second hypothesis is that the extraction kinetics of oolong tea is different from those of other teas, which might result from the extensive drying, physical changes of the leaves (*e.g.* twisting/curling), and oxidation. The molecular composition of the teas changed over time, with observed patterns mainly consistent with continued extraction of molecules as opposed to chemical modifications. While a range of compounds increased in many of the tea types, there were signatures specific to tea type, such as the increase in the relative abundance of coniferyl aldehyde only in oolong tea.

The changes observed are in contrast to the yogurt samples, where chemical alterations over time vary significantly, likely due to the microbial activity. We can detect significant changes in PCoA, molecular networks and heatmaps. In the PCoA, the home ferment inoculated with Kroger yogurt resembled the original starting culture at a molecular level, and it differed from the other home ferments, possibly because it contained a different set of yogurt cultures. Interestingly, significant changes over time are not observed in the heatmap, when we focus our analysis on annotated compounds only (**Figure 4** and **Supplementary Figure 3**), indicating that many of the molecular transformations during fermentation are not yet characterized or that the reference spectra are not present in the available MS library databases. Consistent with the lack of reference spectra in the public databases, the yogurt and milk samples also had the lowest annotation rate at 3.5%. Among the compounds that were annotated we found a broad range of compounds (**Supplementary Figure 6**), including food additives and sugars, which are also found in other milk types within publically available datasets on GNPS, such as breast milk.

One of the questions of interest was whether the different origins of tomatoes could be distinguished on a chemical level. Expectedly, the processed tomatoes (canned and sun-dried) were significantly different from fresh ones. Many molecules including added oils, sugars and preservatives explain these differences. However, differences between fresh fruit are also noticeable in the PCoA data. The private garden-grown tomatoes were used as an ideal case scenario ‑ these fruits were naturally grown, ripened on the vine and have not been treated with any pesticides/herbicides. In PCoA space, farmer’s market tomatoes most closely resemble home-grown ones, while the store-bought tomatoes were all similar to each other. Also, different brands could be distinguished. It is likely that the close similarity of garden and farmer’s market tomatoes results from similar treatment where the fruits are ripened on the vine and collected and sold without any processing (this is known for the garden tomatoes and presumed for the farmer’s market ones). Conversely, the store-bought tomatoes are collected at an early stage, often not fully ripened for ease of transportation, transported over long distances and treated with exogenous ethylene (depending on the supplier). This appears to have a more significant effect on the chemical composition of tomatoes than the “organic” designation.

Another question of interest was whether organic designation and the addition of an antibiotic would impact meat spoiling over time. While the largest difference was that of beef versus turkey, there are some minor trends that can be observed over time in the PCoA plots. Because there was a large within intra-day sample variation, specific major trends could not be detected with respect to organic or antibiotic addition. The data as well as visual inspection of the meat indicate that there were non-uniform chemical transformations, possibly related to the surface area and exposure to air. When the data from each time point is merged, as done with molecular networking, and assessed for presence and absence of spectra, there are hints that there are few low intensity molecular clusters, including oleoyl-taurine and acetyl-carnitine, that change in both the molecular networking data and in a time-dependent manner (**Supplementary Figure 2**). These were different for the different meat classification of organic vs non-organic. Similarly the effect of tetracycline is observed in both the turkey and beef, but only affects few molecules within the 5 day experiment. We expect that these observations are just the tip of the iceberg that warrant further investigation. Future studies can further utilize the mass spectral molecular networking data, with the ability to propagate annotations across a network, to better understand the effect of time on spoilage.

Finally, we assessed the effect of roasting type of coffee across solid “ground” coffee as well as liquid “brewed” coffee. The largest molecular changes were observed between the liquid and solid samples, whereas, comparatively the roasting type only displayed minor molecular changes. This finding suggests that extraction method has a larger effect on the molecular composition of coffee than processing type such as roasting. Alternatively, molecular changes induced by roasting might be predominantly observed in volatile components, not assessed in this study. Indeed, changes in smell between the different roasting types could be readily perceived. Further analyses which address aromatics, such as GC-MS, would be needed to confirm this hypothesis.

In summary we have created five unique data sets that enable the molecular assessment of five common foods and beverages, which are connected by frequently used handling and processing practices. We show that the combination of molecular networking, and multivariate statistical methods such as PCoA and heatmaps and univariate statistics (correlation, significance testing) can be used to explore the molecular composition and the effect of different processing methods, different products and storage conditions relative to all other samples in the study or group. The data sets provided here serve as a reference data set that can continue to be mined. One exemplary feature of the GNPS molecular networking workflow is the search parameter ‘Find Related Datasets’. As exemplified in this study, even the most traditional food types contain a large number of unannotated molecules, therefore, we expect that the increasing deposition of mass spectrometry datasets in the public domain would allow the comparison to other complex mixtures and to narrow down the origin of molecular features. In this spirit the projects are publicly available in GNPS (Wang et al., 2016). Anyone who wishes to continue to learn about these data sets can subscribe to the projects as they will be subjected to living data analysis. Living data is a strategy introduced in Wang et al., 2016 where the data is continuously reanalyzed and updates are provided automatically to all the subscribers of these data sets. As 88-97% of all the signals are currently unannotated, we will, as a community, continue to increase our knowledge about the molecular composition and changes of our food.

## Data and code availability

Data can be accessed via http://gnps.ucsd.edu via accession numbers for Meat (G1): MSV000082423; Tomato (G2): MSV000082391; Coffee (G3): MSV000082386; Milk/yogurt (G4): MSV000082387; Tea (G5): MSV000082388. Metadata are uploaded with each dataset. All jupyter notebooks and scripts used for data pre-processing and analysis are publically available at: https://github.com/DorresteinLaboratory/supplementary-MolecularChangesInFood

Links to networking jobs for GNPS networking jobs: Parameters are precursor and fragment ion tolerance set to 0.1 Da, min matched fragment ions: 4, cosine 0.7; library search min match peaks: 4; run with metadata, attributes assigned; NIST17 included.

G1 meat samples only https://gnps.ucsd.edu/ProteoSAFe/status.jsp?task=1520f7112d9f445384eb743ac4358c21

G2 tomato samples only https://gnps.ucsd.edu/ProteoSAFe/status.jsp?task=cf6e0347de8b48aeb8c700e45b4d3159

G3 coffee samples only https://gnps.ucsd.edu/ProteoSAFe/status.jsp?task=2c83502fefc0469ba79ba70a9461b9b5

G4 yogurt samples only https://gnps.ucsd.edu/ProteoSAFe/status.jsp?task=1f11fb76e15240a893e46e02d9c58cd2

G5 tea samples only https://gnps.ucsd.edu/ProteoSAFe/status.jsp?task=f7569832b0d241f79812446d81cd5ca5

Global: https://gnps.ucsd.edu/ProteoSAFe/status.jsp?task=b881151839574f639ceaf06f9b11e464

Links for jobs performed to obtain the feature MS1 and MS2 linked table for PCoA and heatmaps and statistical analysis: Parameters are precursor and fragment ion tolerance set to 0. 1 Da, 4 min match peaks, cosine 0.7; inputs are the .mgf; .csv from mzMINE and the metadata file.

Global: https://gnps.ucsd.edu/ProteoSAFe/status.jsp?task=25577f11a35c48cdb30dfc005cbd6638

G1 meat: https://gnps.ucsd.edu/ProteoSAFe/status.isp?task=85c7380ca7ee44b3a3ed448c9b4d09fa

G2 tomato: http://gnps.ucsd.edu/ProteoSAFe/status.isp?task=7a4031dd3ee146699ceecef919d7f668

G3 coffee: https://gnps.ucsd.edu/ProteoSAFe/status.isp?task=750cc6fe82dc4732b84b355904cf91d3

G4 yogurt: http://gnps.ucsd.edu/ProteoSAFe/status.isp?task=f9ede96d9e694f9e8b6186991e289c17

G5 tea: https://gnps.ucsd.edu/ProteoSAFe/status.isp?task=80fc6384e52b4fe18e13ebbb3a86b4d8

Link to Clusterapp: http://dorresteinappshub.ucsd.edu:3838/clusterMetaboApp0.9.1/

## Acknowledgments

The result of this work was a part of a hands-on mass spectrometry course at UCSD called “System Wide Mass Spectrometry”. The authors were all participants of this course. We further acknowledge NIH number P41 GM103484, and NIH Grant GMS10RR029121 and Bruker for the shared instrumentation infrastructure that enabled this work and the UCSD Center for Microbiome Innovation.

## Supplementary Information

**Movie S1.**
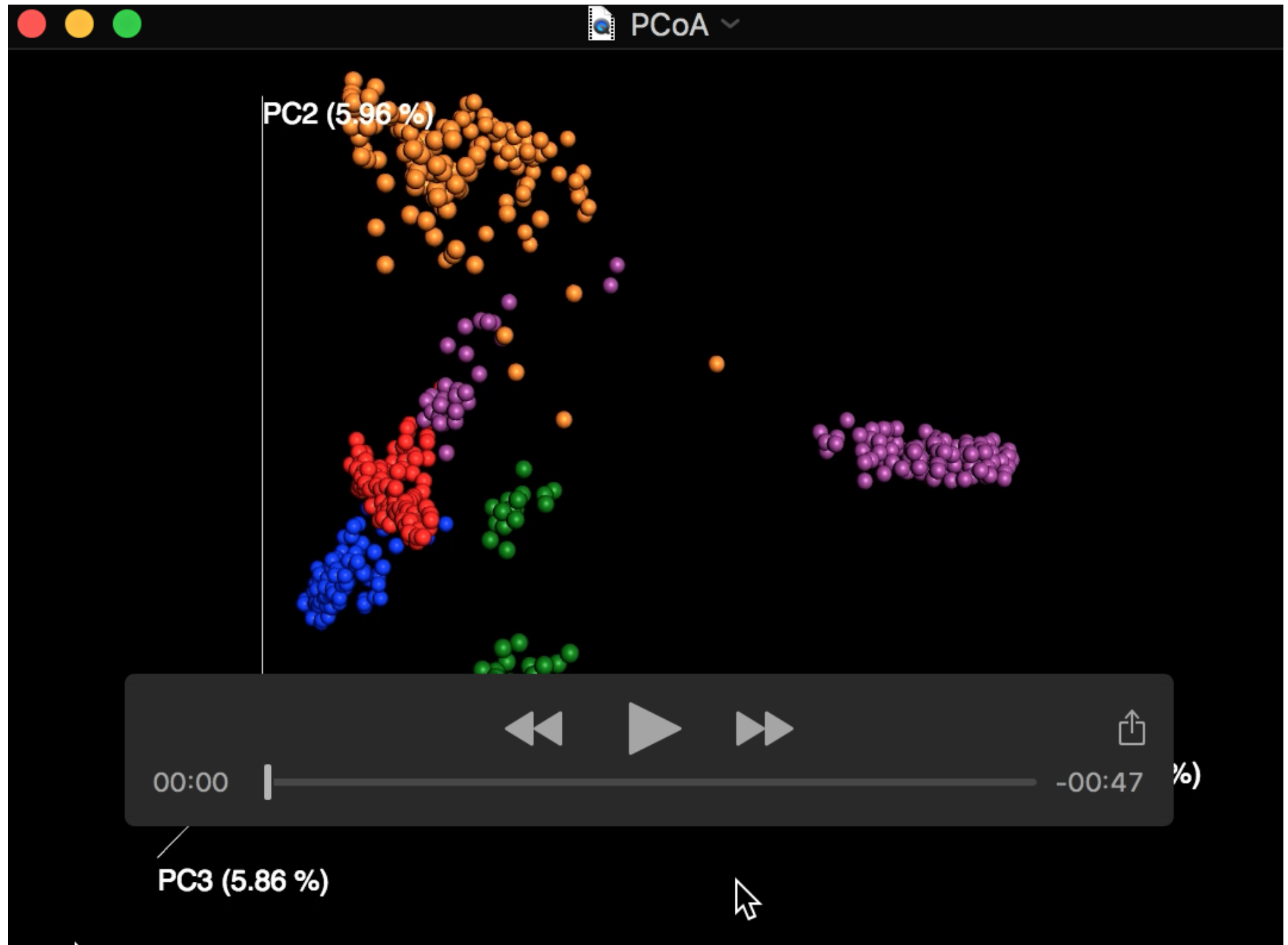
PCoA of all the samples for 3D visualization.

**Supplementary Figure 1.**
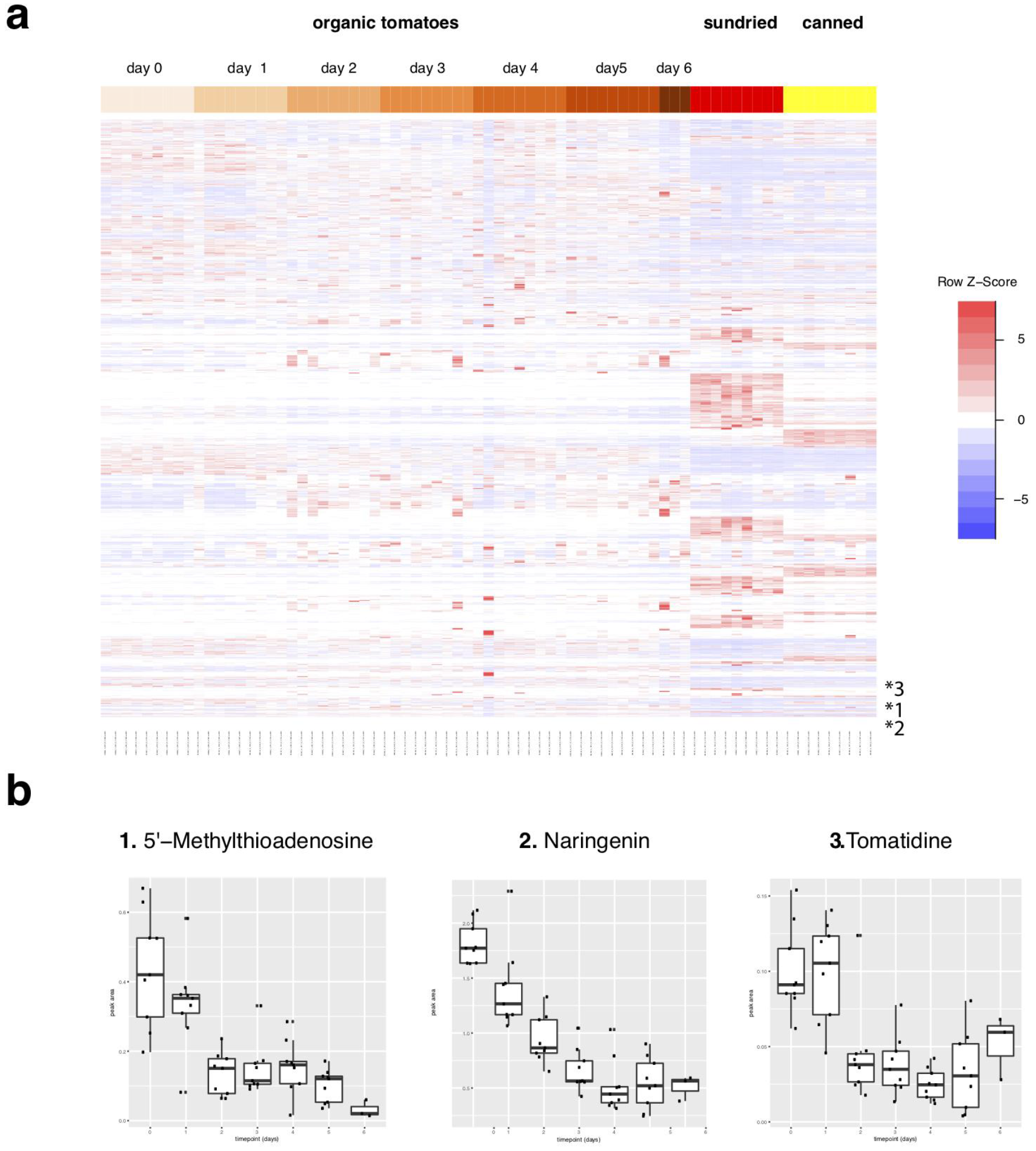
Heatmap showing tomato metabolites changing over time and tomato type. **a.** Heatmap of organic fresh tomato, sundried, and canned tomato metabolites changing over time **b.** Selected metabolites found within the organic tomatoes significantly decrease in their relative intensity over time upon ripening at room temperature, including 5’Methylthioadenosine.

**Supplementary Figure 2.**
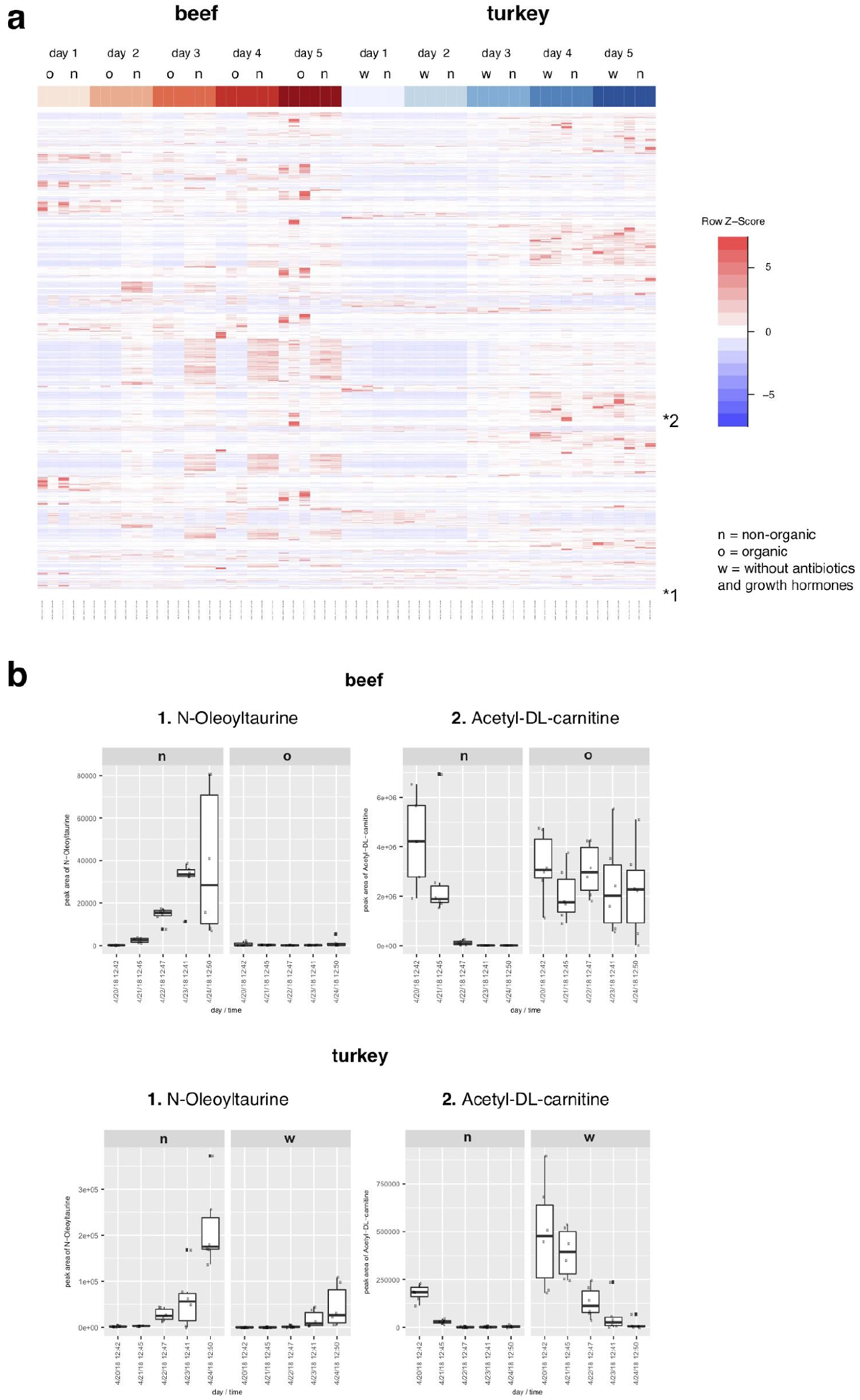
Heatmap showing metabolites changing from meat samples. **a.** Heatmap of beef (organic and non-organic) and turkey (with and without antibiotic/growth hormones). **b.** Selected metabolites found within the organic or non-organic meat significantly changed in their relative intensity over the period of 5 days. o = organic and no tetracycline, n = non-organic and no tetracycline, w = with tetracycline.

**Supplementary Figure 3.**
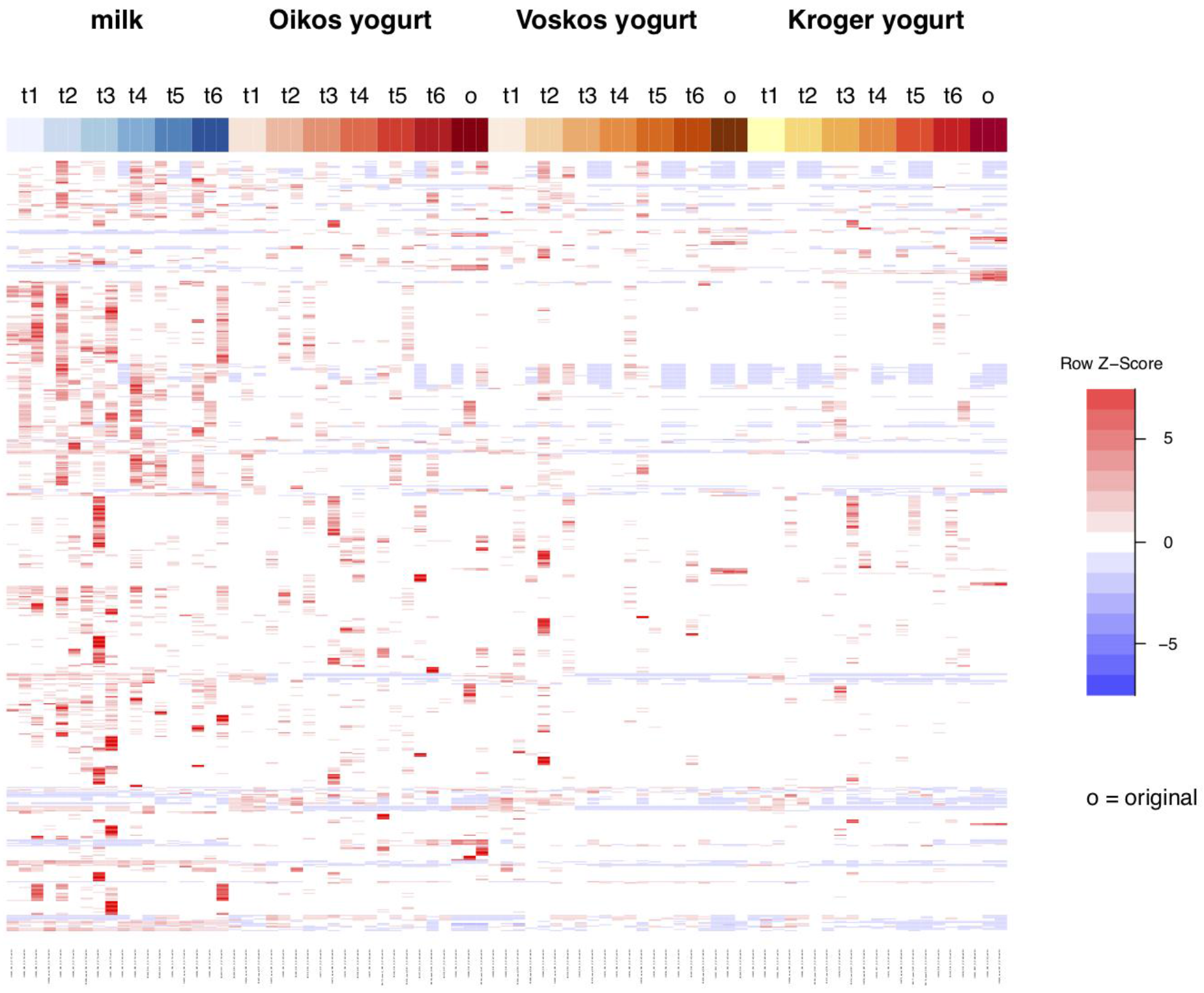
Heatmap showing annotated metabolites changing during the fermentation process from milk to yogurt across different yogurt brands revealing a general lack of trends. Metabolite annotation was performed through mass spectral molecular networking and spectral matching to reference spectra.

**Supplementary Figure 4.**
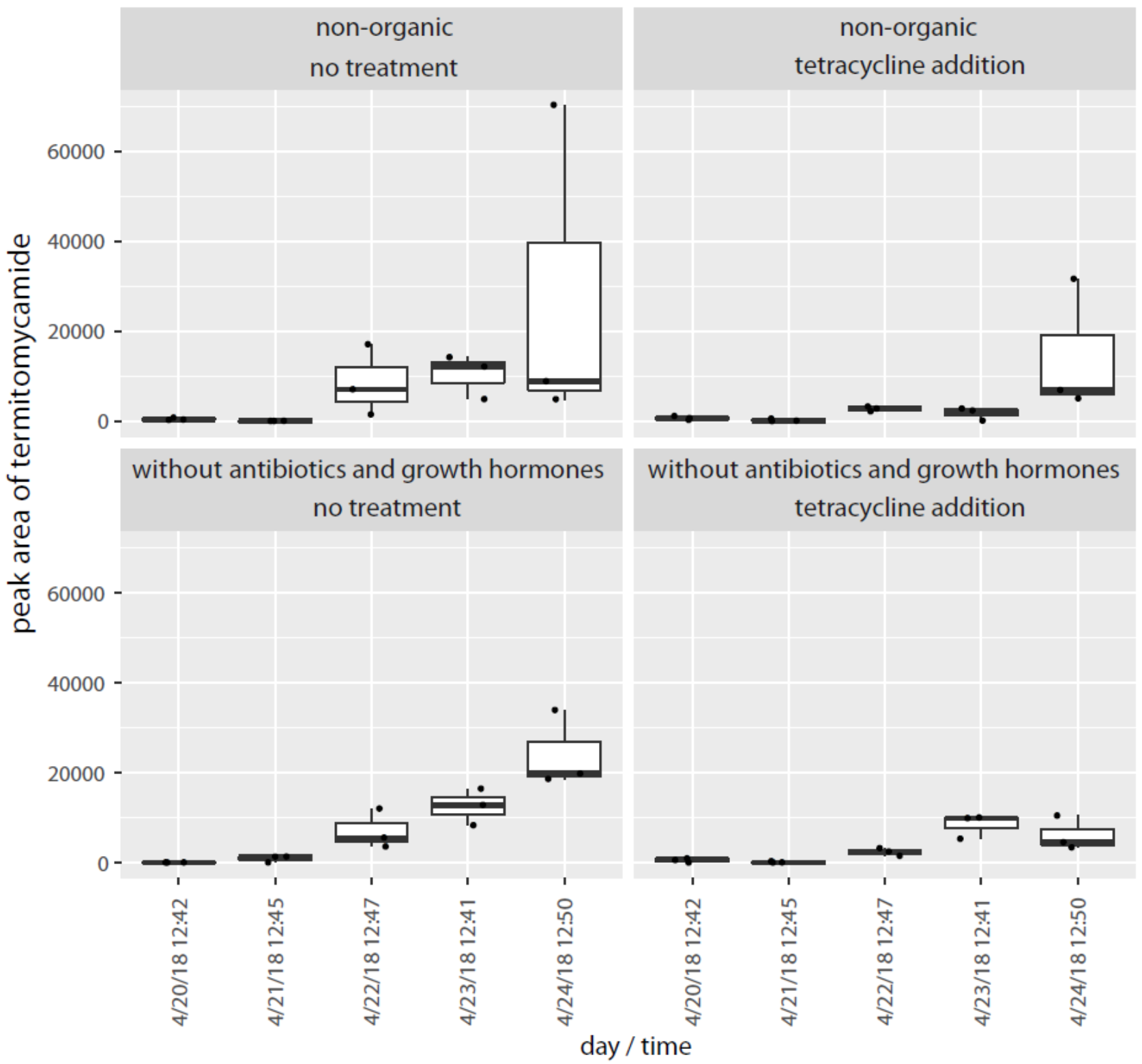
The kinetics of termitomycamide abundance over time in turkey samples. Top is non-organic turkey (Kroger brand). Bottom is turkey grown without antibiotics and growth hormones (Empire brand).

**Supplementary Figure 5.**
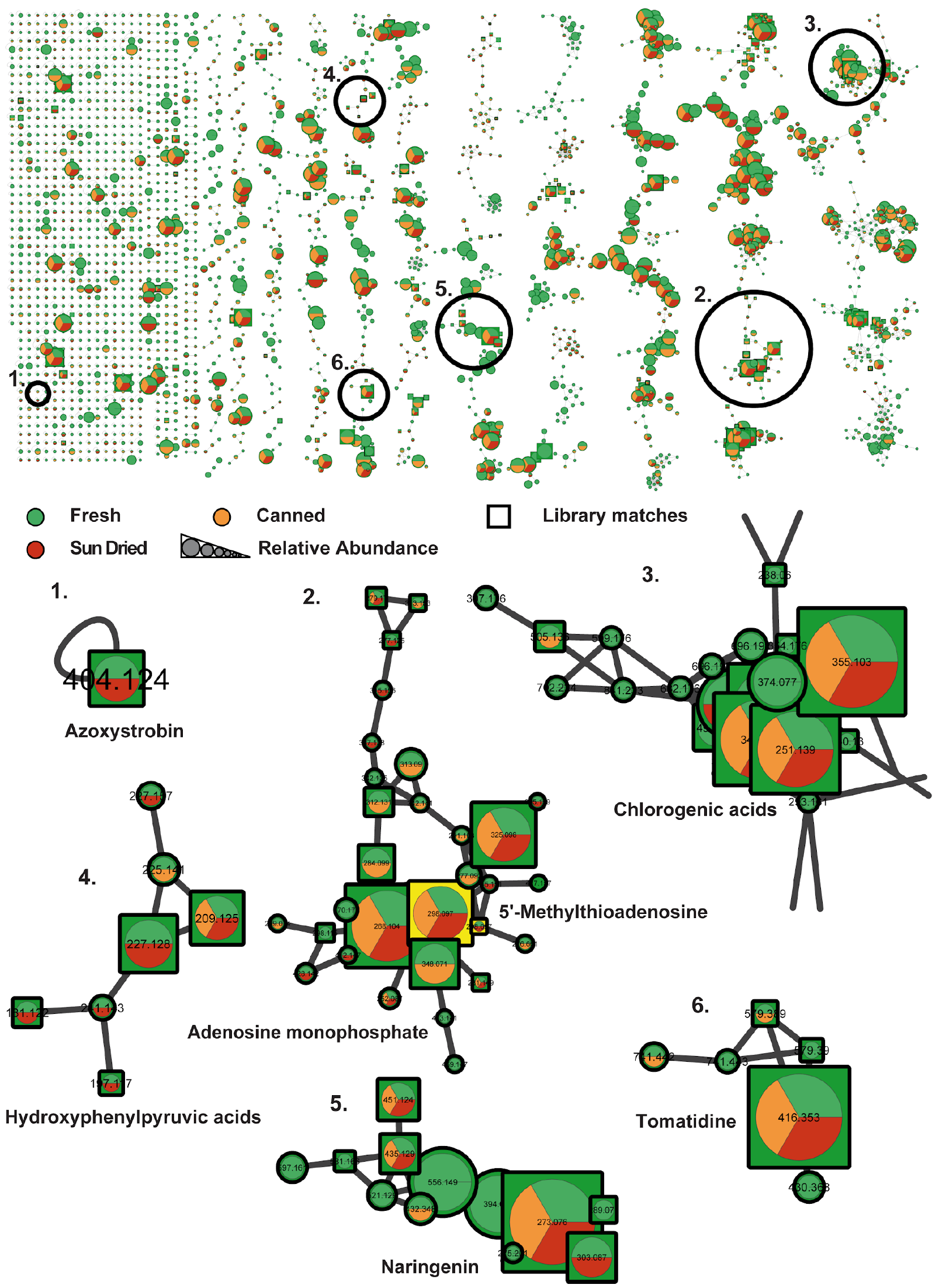
The complete tomato molecular network from **Figure 5a**. Molecular network shows a global overview of the spectral similarity of all MS/MS spectra from this dataset. Below the global network shown in the upper panel numbered zoomed regions are shown. Binary presence of MS/MS spectra in different subsets of the samples are indicated through the color coding of pie charts. Node sizes indicated relative precursor abundance and selected library identifications are annotated in the figure or shown through squared node shape.

**Supplementary Figure 6.**
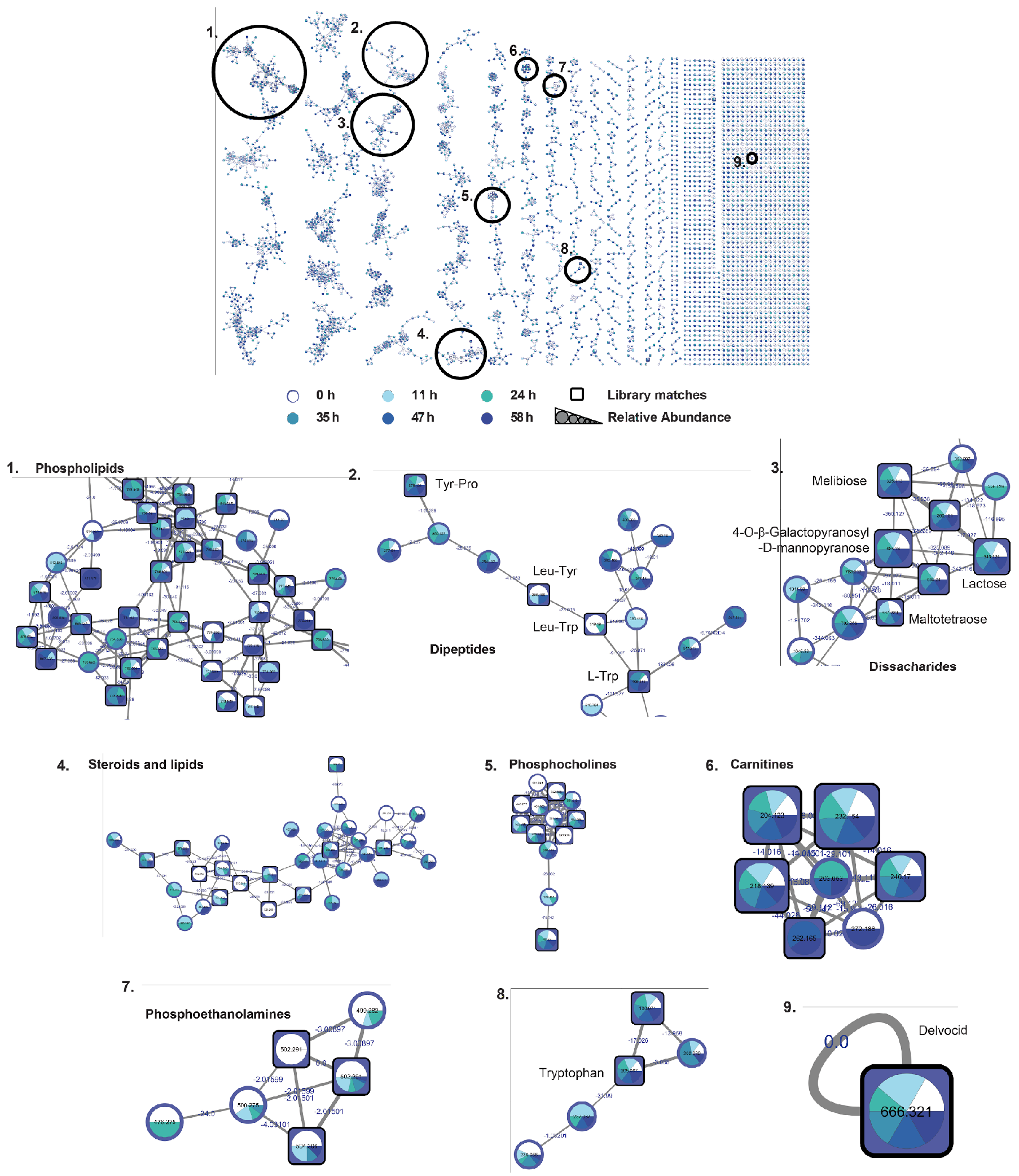
The complete milk to yogurt molecular network from **Figure 5b**. Molecular network shows a global overview of the spectral similarity of all MS/MS spectra from this dataset. Numbered amplified regions in the global network are shown. Contribution of MS/MS spectra from different fermentation time points are indicated by the color coding of pie charts. Node sizes indicates relative precursor abundance. Library identifications are represented with squared node shapes.

**Supplementary Figure 7.**
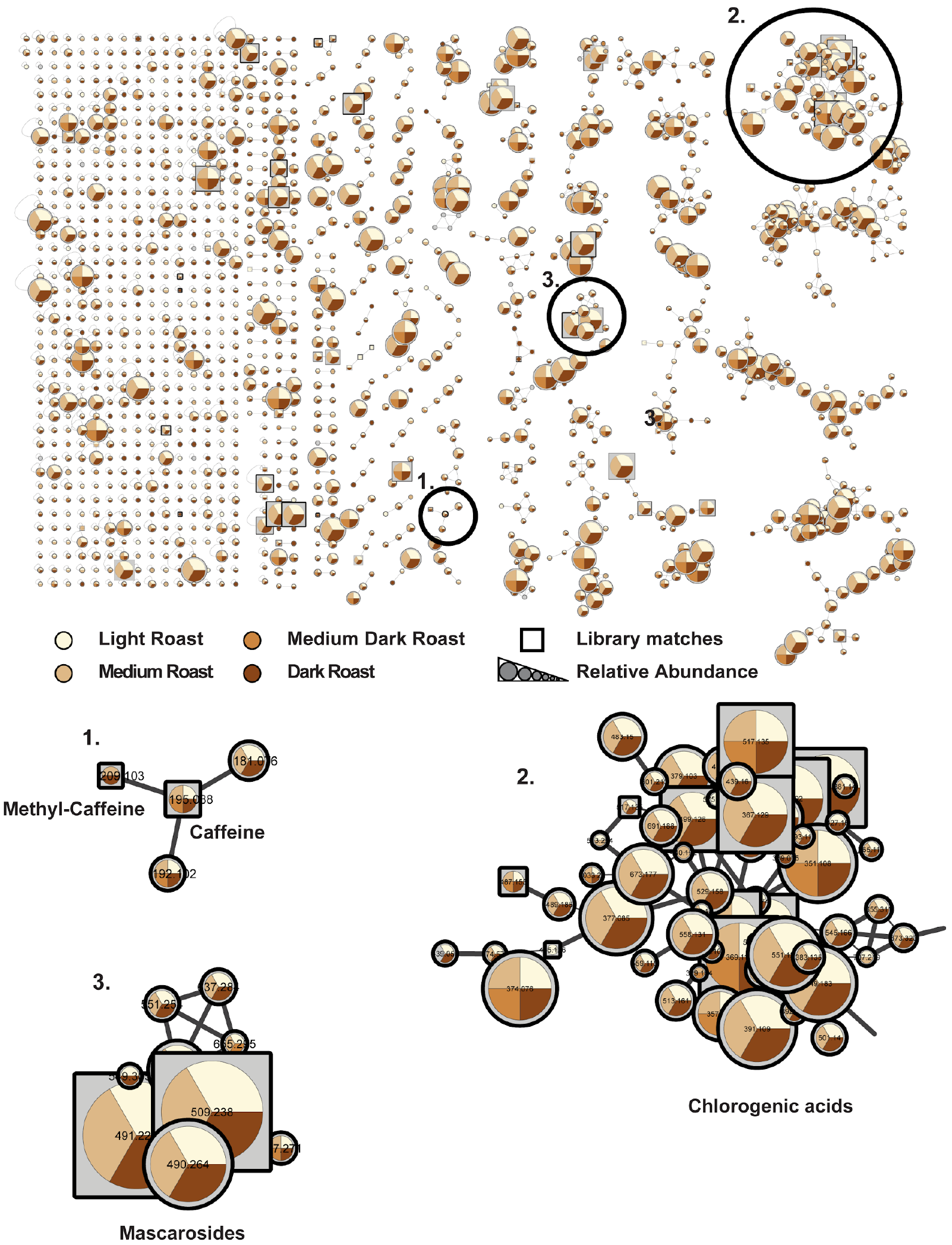
The complete coffee molecular network from **Figure 5c**. The Molecular network shows a global overview of the spectral similarity of all MS/MS spectra from this dataset. Below the global network shown in the upper panel numbered zoomed regions are shown. Binary presence of MS/MS spectra in different roast type subsets of the samples are indicated through the color coding of pie charts. Node sizes indicate relative precursor abundance and selected library identifications are annotated in the figure or shown through squared nodeshape.

**Supplementary Figure 8.**
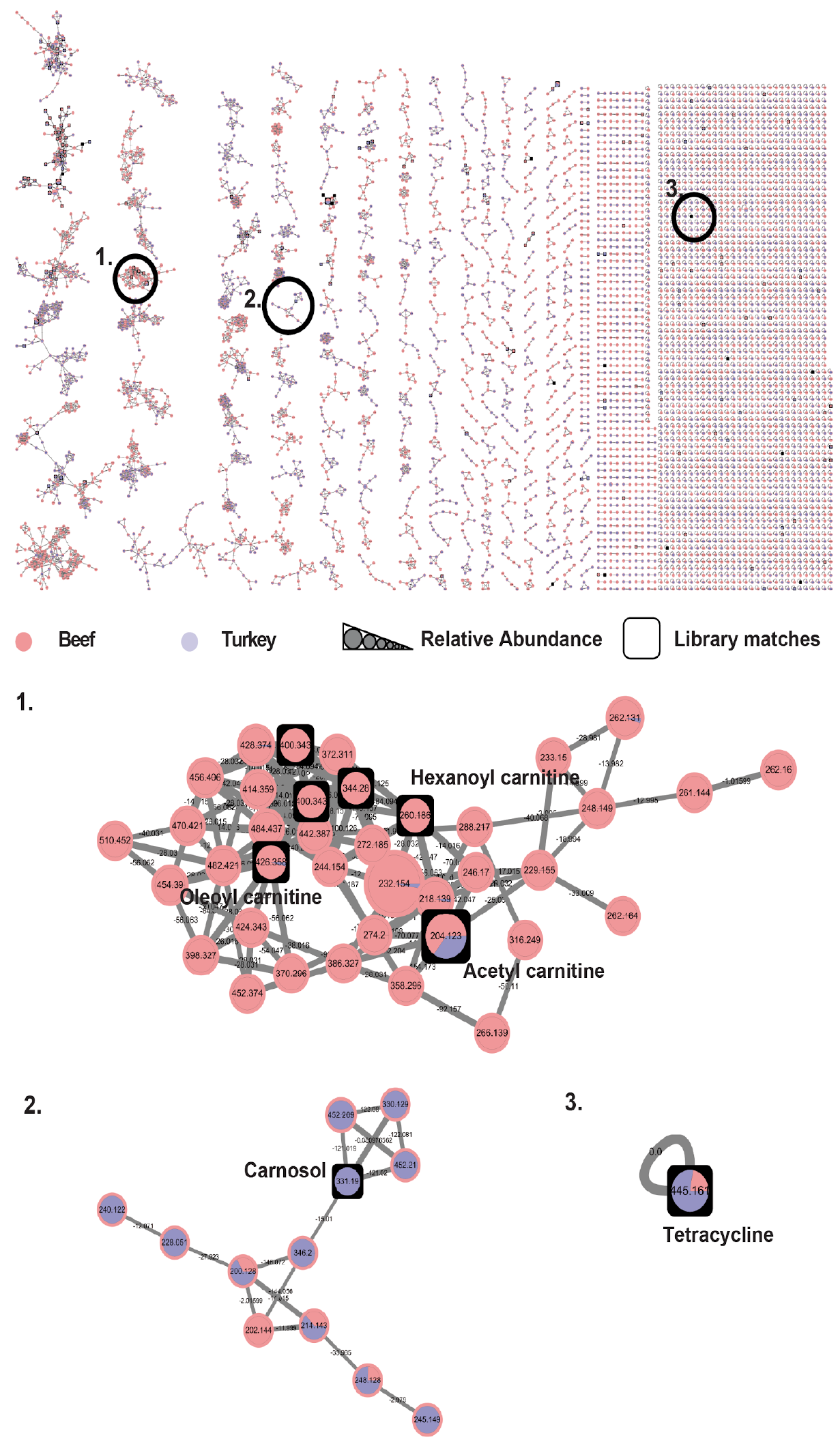
The complete meat molecular network from **Figure 6a**. Molecular network shows a global overview of the spectral similarity of all MS/MS spectra from this dataset. Below the global network shown in the upper panel numbered zoomed regions are shown. Spectral counts of MS/MS spectra in different meat type subsets of the samples are indicated through the color coding and ratio of pie charts. Node sizes indicated relative precursor abundance and selected library identifications are annotated in the figure or shown through squared node shape.

**Supplementary Figure 9.**
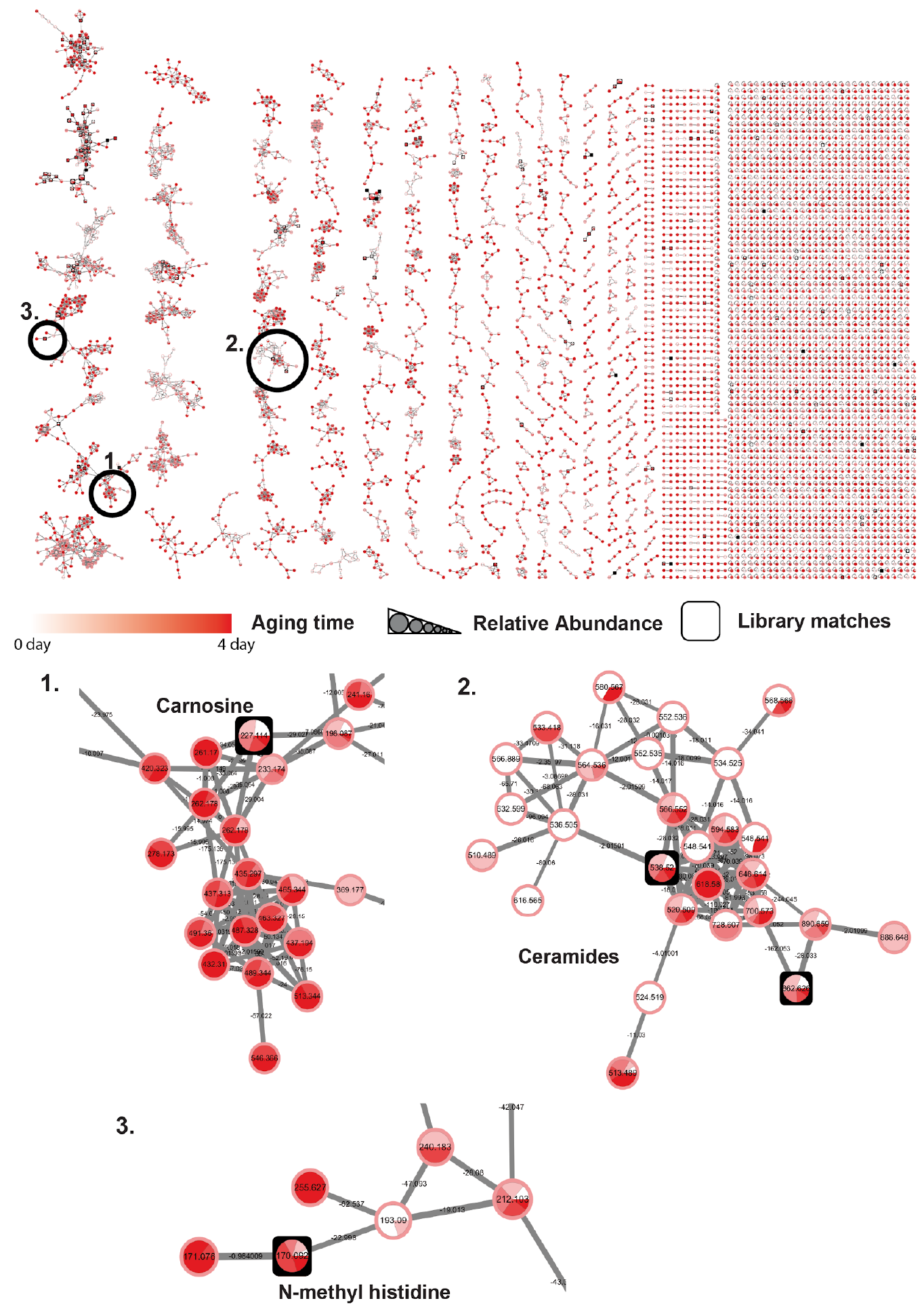
The complete meat molecular network from **Figure 6b**. Molecular network shows a global overview of the spectral similarity of all MS/MS spectra from this dataset. Below the global network shown in the upper panel numbered zoomend regions are shown. Spectral counts of MS/MS spectra in different subsets of the samples are indicated through the color coding and ratio of pie charts. Node sizes indicated relative precursor abundance and selected library identifications are annotated in the figure or shown by a squared node shape.

**Supplementary Figure 10.**
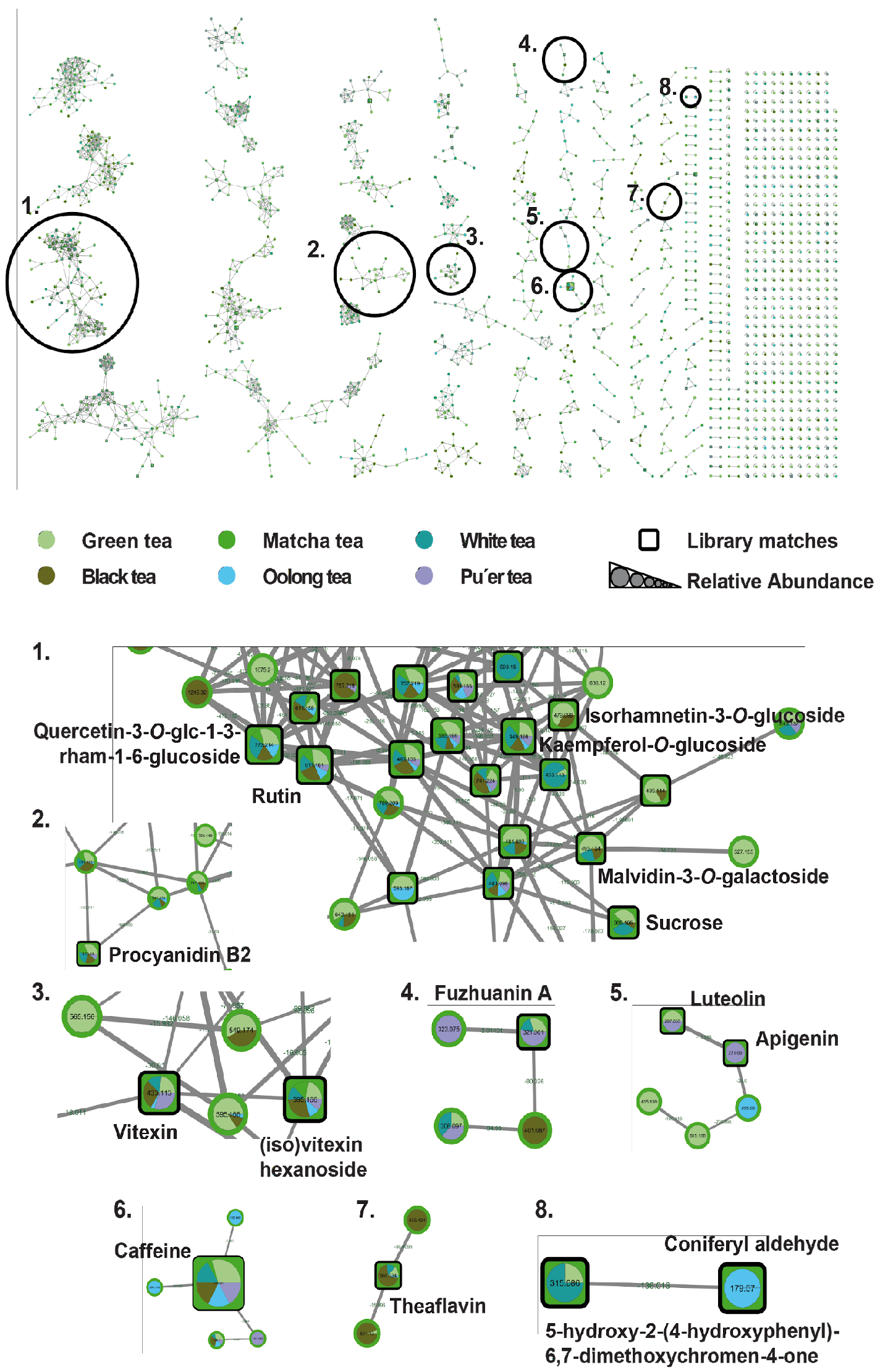
The completer tea molecular network from **Figure 6c**. Molecular network shows a global overview of the spectral similarity of all MS/MS spectra from this dataset. Amplified regions indicated by numbers in the global network (up) are shown. Contribution of MS/MS spectra in different tea types are indicated through the color coding of pie charts. Node sizes indicates relative precursor abundance. Library identifications are represented with squared node shapes.

**Supplementary Figure 11.**
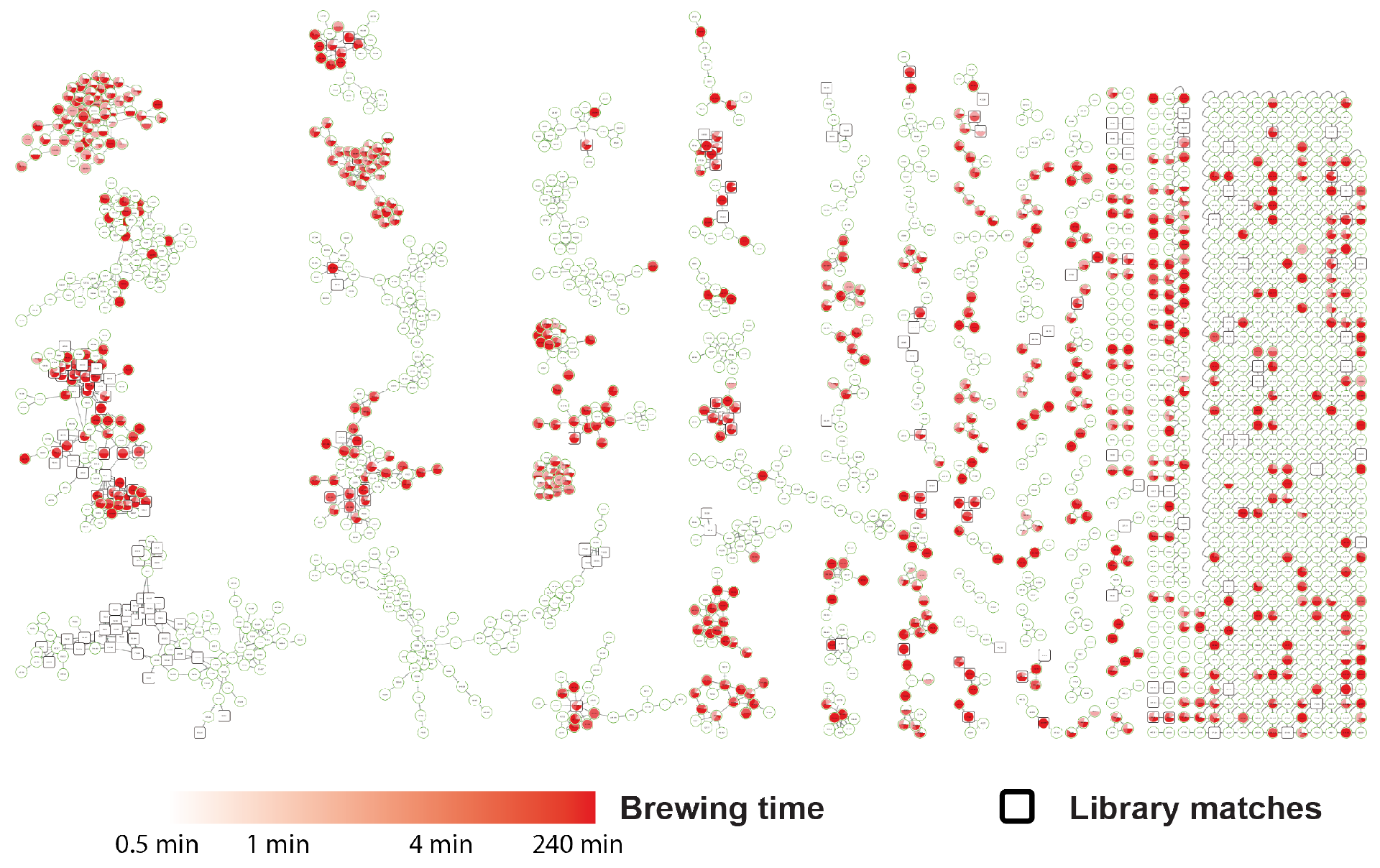
Molecular network of brewing time for tea, the network is the same as **Supplementary Figure 10** but with the metadata of time used to color it.

**Supplementary Figure 12.**
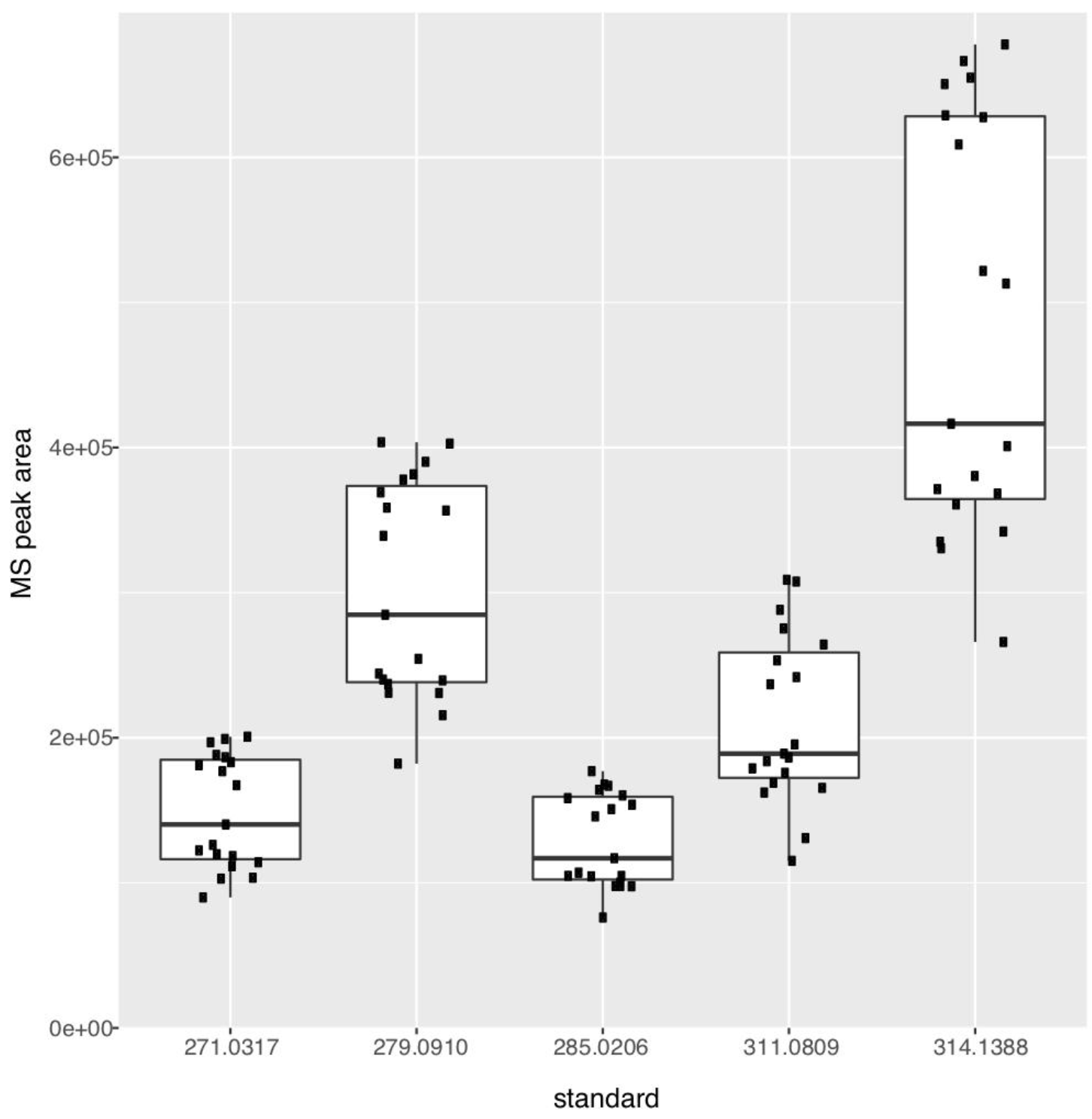
Plot of standards in standard mix, injected on average every 50 samples. Shows instrument consistency across the time course of all sample groups run on the QTOF mass spectrometer. The minor grouping separation correspond to instrument maintenance (cleaning the MS source and changing the guard cartridge). Standard mix was Sulfamethazine [*m/z* 279.0910 (RSD 24.95)], Sulfamethizole [*m/z* 271.0317 (RSD 26.08)] Sulfachloropyridazine [*m/z* 285.0206 (RSD 24.94)], Sulfadimethoxine [311.0809 (RSD 27.55)], Coumarin-314 [314.1388 (RSD 29.61)]. This means that variations in our experiments are larger than the RSDs, which we observe here.

## References (prefer 40 max and if we need more ignore the 40 cut-off but needs to be alphabetical by first author

Aksenov, A. A., da Silva, R., Knight, R., Lopes, N. P., & Dorrestein, P. C. (2017). Global chemical analysis of biology by mass spectrometry. Nature Reviews Chemistry, 1, 0054.

Blaženović, I., Kind, T., Ji, J., & Fiehn, O. (2018). Software tools and approaches for compound identification of LC-MS/MS data in metabolomics. Metabolites, 8(2), 31.

Borges, R. M., Taujale, R., de Souza, J. S., de Andrade Bezerra, T., Silva, E. L. E., Herzog, R., Ponce, F. V., Wolfender, J. L., & Edison, A. S. (2018). Dereplication of plant phenolics using a mass-spectrometry database independent method. Phytochemical Analysis, (in press) doi:10.1002/pca.2773.

Branen, A. L., Davidson, P. M., Salminen, S., & Thorngate, J. (2001). Food Additives. New York: CRC Press. pp. 599–600. ISBN 9780824741709.

Caporaso, J. G., Kuczynski, J., Stombaugh, J., Bittinger, K., Bushman, F. D., Costello, E. K., Fierer, N., Peña, A. G., Goodrich, J. K., Gordon, J. I., Huttley, G. A., Kelley, S. T., Knights, D., Koenig, J. E., Ley, R. E., Lozupone, C. A., McDonald, D., Muegge, B. D., Pirrung, M., Reeder, J., Sevinsky, J. R., Turnbaugh, P. J., Walters, W. A., Widmann, J., Yatsunenko, T., Zaneveld, J., & Knight, R. (2010). QIIME allows analysis of high-throughput community sequencing data. Nature Methods, 7(5), 335–336.

Casida, J. E. & Durkin, K. A. (2017). Pesticide chemical research in toxicology: lessons from Nature. Chemical Research in Toxicology, 30(1), 94–104.

Choi, J. H., Maeda, K., Nagai, K., Harada, E., Kawade, M., Hirai, H., & Kawagishi, H. (2010). Termitomycamides A to E, fatty acid amides isolated from the mushroom Termitomyces titanicus, suppress endoplasmic reticulum stress. Organic Letters, 12(21), 5012–5015.

Clifford, M. N., Jaganath, I. B., Ludwig, I. A., & Crozier, A. (2017). Chlorogenic acids and the acyl-quinic acids: discovery, biosynthesis, bioavailability and bioactivity. Natural Product Reports, 34(12), 1391–1421.

Cuyckens, F. & Claeys, M. (2004). Mass spectrometry in the structural analysis of flavonoids. Journal of Mass Spectrometry, 39(1), 1–15.

Ejugi, B. A., Valkenborg, D., Baggerman, G., Vanaerschot, M, Witters, E., Dujardin, J. C., Burzykowski, T., & Berg, M. (2013). Evaluation of normalization methods to pave the way towards large-scale LC-MS-based metabolomics profiling experiments. Omics, 17(9), 473—485.

Floros, D. J., Petras, D., Kapono, C. A., Melnik, A. V., Ling, T.-J., Knight, R., & Dorrestein, P. C. (2017). Mass spectrometry based molecular 3D-cartography of plant metabolites. Frontiers in Plant Science, 8, 429.

Ge, Y. W., Zhu, S., Yoshimatsu, K., & Komatsu, K. (2017). MS/MS similarity networking accelerated target profiling of triterpene saponins in Eleutherococcus senticosus leaves. Food Chemistry, 227, 444–452.

Giorio, C, Safer, A., Sánchez-Bayo, F., Tapparo, A., Lentola, A., Girolami, V., van Lexmond, M. B., & Bonmatin, J.M. (2017). An update of the Worldwide Integrated Assessment (WIA) on systemic insecticides. Part 1: new molecules, metabolism, fate, and transport. Environmental Science and Pollution Research. (in press) doi:10.1007/s11356-017-0394-3.

Granados-Chinchilla, F. & Rodríguez, C. (2017). Tetracyclines in food and feedingstuffs: From regulation to analytical methods, bacterial resistance, and environmental and health implications. Journal of Analytical Methods in Chemistry, 2017, 1315497.

Islam, M. T., Tabrez, S., Jabir, N. R., Ali, M., Kamal, M. A., da Silva Araujo, L., De Oliveira Santos, J. V., Da Mata, A. M. O. F., De Aguiar, R. P. S., & de Carvalho Melo Cavalcante, A. A. (2018). An insight on the therapeutic potential of major coffee components. Current Drug Metabolism, (in press) doi:10.2174/1389200219666180302154551.

Karpinska, J., Świsłocka, R., & Lewandowski, W. (2017). A mystery of a cup of coffee; an insight look by chemist. Biofactors, 43(5), 621–632.

Luo, Z. M., Du, H. X., Li, L. X., An, M. Q., Zhang, Z. Z., Wan, X. C., Bao, G. H., Zhang, L., & Ling, T. J. (2013). Fuzhuanins A and B: the B-ring fission lactones of flavan-3-ols from Fuzhuan brick-tea. Journal of Agricultural and Food Chemistry, 61(28), 6982–6990.

Marshall, J. W., Schmitt-Kopplin, P., Schuetz, N., Moritz, F., Roullier-Gall, C., Uhl, J., Colyer, A., Jones, L. L., Rychlik, M., & Taylor, A. J. (2018). Monitoring chemical changes during food sterilisation using ultrahigh resolution mass spectrometry. Food Chemistry, 242, 316–322.

Naveed, M., BiBi, J., Kamboh, A. A., Suheryani, I., Kakar, I., Fazlani, S. A., FangFang, X., Kalhoro, S. A., Yunjuan, L., Kakar, M. U., Abd El-Hack, M. E., Noreldin, A. E., Zhixiang, S., LiXia, C., & XiaoHui, Z. (2018a). Pharmacological values and therapeutic properties of black tea (Camellia sinensis): A comprehensive overview. Biomedicine & Pharmacotherapy, 100, 521–531.

Naveed, M., Hejazi, V., Abbas, M., Kamboh, A. A., Khan, G. J., Shumzaid, M., Ahmad, F., Babazadeh, D., FangFang, X., Modarresi-Ghazani, F., WenHua, L., & XiaoHui, Z. (2018b). Chlorogenic acid (CGA): A pharmacological review and call for further research. Biomedicine & Pharmacotherapy, 97:67–74.

Nguyen, D. D., Wu, C. H., Moree, W. J., Lamsa, A., Medema, M. H., Zhao, X., Gavilan, R. G., Aparicio, M., Atencio, L., Jackson, C., Ballesteros, J., Sanchez, J., Watrous, J. D., Phelan, V. V., van de Wiel, C., Kersten, R. D., Mehnaz, S., De Mot, R., Shank, E. A., Charusanti, P., Nagarajan, H., Duggan, B. M., Moore, B. S., Bandeira, N., Palsson, B. O., Pogliano, K., Gutiérrez, M., & Dorrestein, P. C. (2013). MS/MS networking guided analysis of molecule and gene cluster families. Proceedings of the National Academy of Sciences of the United States of America, 110(28), E2611–E2620.

North, J. A., Miller, A. R., Wildenthal, J. A., Young, S. J., & Tabita, F. R. (2017). Microbial pathway for anaerobic 5′-methylthioadenosine metabolism coupled to ethylene formation. Proceedings of the National Academy of Sciences of the United States of America, 114(48), E10455–E10464.

Pastoriza, S., Mesías, M., Cabrera, C., & Rufián-Henares, J. A. (2017). Healthy properties of green and white teas: an update. Food & Functions, 8(8), 2650–2662.

Scalbert, A., Brennan, L., Manach, C., Andres-Lacueva, C., Dragsted, L. O., Draper, J., Rappaport, S. M., van der Hooft, J. J. J., & Wishart, D. S. (2014). The food metabolome: a window over dietary exposure. American Journal of Clinical Nutrition, 99(6), 1286–1308.

Scheubert, K., Hufsky, F., Petras, D., Wang, M., Nothias, L.-F., Dührkop, K., Bandeira, N., Dorrestein, P. C., & Böcker, S. (2017). Significance estimation for large scale metabolomics annotations by spectral matching. Nature Communication, 8(1), 1494.

Shu, Y., Liu, J. Q., Peng, X. R., Wan, L. S., Zhou, L., Zhang, T., & Qiu, M. H. (2014). Characterization of diterpenoid glucosides in roasted puer coffee beans. Journal of Agricultural and Food Chemistry, 62(12), 2631–2637.

Souard, F., Delporte, C., Stoffelen, P., Thévenot, E. A., Noret, N., Dauvergne, B., Kauffmann, J. M., Van Antwerpen, P., & Stévigny, C. (2018). Metabolomics fingerprint of coffee species determined by untargeted-profiling study using LC-HRMS. Food Chemistry, 245, 603–612.

Sumner, L. W., Amberg, A., Barrett, D., Beale, M. H., Beger, R., Daykin, C. A., Fan, T. W.-M., Fiehn, O., Goodacre, R., Griffin, J. L., Hankemeier, T., Hardy, N., Harnly, J., Higashi, R., Kopka, J., Lane, A. N., Lindon, J. C., Marriott, P., Nicholls, A. W., Reily, M. D., Thaden, J. J., & Viant, M. R. (2007). Proposed minimum reporting standards for chemical analysis Chemical Analysis Working Group (CAWG) Metabolomics Standards Initiative (MSI). Metabolomics, 3(3), 211–221.

Tajik, N., Tajik, M., Mack, I., & Enck, P. (2017). The potential effects of chlorogenic acid, the main phenolic components in coffee, on health: a comprehensive review of the literature. European Journal of Nutrition, 56(7), 2215–2244.

Turman, M. V., Kingsley, P. J., Rouzer, C. A., Cravatt, B. F., & Marnett, L. J. (2008). Oxidative metabolism of a fatty acid amide hydrolase-regulated lipid, arachidonoyltaurine. Biochemistry, 47(12), 3917–3925.

van der Hooft, J. J. J., Akermi, M., Ünlü, F. Y., Mihaleva, V., Roldan, V. G., Bino, R. J., de Vos, R. C., & Vervoort, J. (2012). Structural annotation and elucidation of conjugated phenolic compounds in black, green, and white tea extracts. Journal of Agricultural and Food Chemistry, 60(36), 8841–8850.

van der Hooft, J. J. J., Vervoort, J., Bino, R. J., & de Vos, R. C. H. (2012). Spectral trees as a robust annotation tool in LC-MS based metabolomics. Metabolomics, 8(4), 691–703.

Vázquez-Baeza, Y., Pirrung, M., Gonzalez, A., & Knight, R. (2013). EMPeror: a tool for visualizing high-throughput microbial community data. Gigascience, 2(1), 16.

Wang, M., Carver, J. J., Phelan, V. V., Sanchez, L. M., Garg, N., Peng, Y., Nguyen, D. D., Watrous, J., Kapono, C. A., Luzzatto-Knaan, T., Porto, C., Bouslimani, A., Melnik, A. V., Meehan, M. J., Liu, W. T., Crüsemann, M., Boudreau, P. D., Esquenazi, E., Sandoval-Calderón, M., Kersten, R. D., Pace, L. A., Quinn, R. A., Duncan, K. R., Hsu, C. C., Floros, D. J., Gavilan, R. G., Kleigrewe, K., Northen, T., Dutton, R. J., Parrot, D., Carlson, E. E., Aigle, B., Michelsen, C. F., Jelsbak, L., Sohlenkamp, C., Pevzner, P., Edlund, A., McLean, J., Piel, J., Murphy, B. T., Gerwick, L., Liaw, C. C., Yang, Y.-L., Humpf, H. U., Maansson, M., Keyzers, R. A., Sims, A. C., Johnson, A. R., Sidebottom, A. M., Sedio, B.E., Klitgaard, A., Larson, C. B., Boya P, C. A., Torres-Mendoza, D., Gonzalez, D. J., Silva, D. B., Marques, L. M., Demarque, D. P., Pociute, E., O’Neill, E. C., Briand, E., Helfrich, E. J. N., Granatosky, E. A., Glukhov, E., Ryffel, F., Houson, H., Mohimani, H., Kharbush, J. J., Zeng, Y., Vorholt, J. A., Kurita, K. L., Charusanti, P., McPhail, K. L., Nielsen, K. F., Vuong, L., Elfeki, M., Traxler, M. F., Engene, N., Koyama, N., Vining, O. B., Baric, R., Silva, R. R., Mascuch, S. J., Tomasi, S., Jenkins, S., Macherla, V., Hoffman, T., Agarwal, V., Williams, P. G., Dai, J., Neupane, R., Gurr, J., Rodríguez, A. M. C., Lamsa, A., Zhang, C., Dorrestein, K., Duggan, B. M., Almaliti, J., Allard, P. M., Phapale, P., Nothias, L.-F., Alexandrov, T., Litaudon, M., Wolfender, J. L., Kyle, J. E., Metz, T. O., Peryea, T., Nguyen, D. T., VanLeer, D., Shinn, P., Jadhav, A., Müller, R., Waters, K. M., Shi, W., Liu, X., Zhang, L., Knight, R., Jensen, P. R., Palsson, B. O., Pogliano, K., Linington, R. G., Gutiérrez, M., Lopes, N. P., Gerwick, W. H., Moore, B. S., Dorrestein, P. C., & Bandeira, N. (2016). Sharing and community curation of mass spectrometry data with Global Natural Products Social Molecular Networking. Nature Biotechnology, 34(8), 828–837.

Watrous, J., Roach, P., Alexandrov, T., Heath, B. S., Yang, J. Y., Kersten, R. D., van der Voort, M., Pogliano, K., Gross, H., Raaijmakers, J. M., Moore, B. S., Laskin, J., Bandeira, N., & Dorrestein, P. C. (2012). Mass spectral molecular networking of living microbial colonies. Proceedings of the National Academy of Sciences of the United States of America, 109(26), E1743–E1752.

Yang, J.-Y., Sanchez, L. M., Rath, C. M., Liu, X., Boudreau, P. D., Bruns, N., Glukhov, E., Wodtke, A., de Felicio, R., Fenner, A., Wong, W. R., Linington, R. G., Zhang, L., Debonsi, H. M., Gerwick, W. H., & Dorrestein, P. C. (2013). Molecular networking as a dereplication strategy. Journal of Natural Products, 76(9), 1686–1699.

Yoshimura, Y., Goto-Inoue, N., Moriyama, T., & Zaima, N. (2016). Significant advancement of mass spectrometry imaging for food chemistry. Food Chemistry, 210, 200–211.

Zhang, S., Yang, C., Idehen, E., Shi, L., Lv, L., & Sang, S. (2018). Novel theaflavin-type chlorogenic acid derivatives identified in black tea. Journal of Agricultural and Food Chemistry, 66(13), 3402–3407.

